# A collection of diverse bacteriophages for biocontrol of ESBL- and AmpC-β-lactamase-producing *E. coli*

**DOI:** 10.1101/2023.09.14.557699

**Authors:** Amira R. Vitt, Anders Nørgaard Sørensen, Martin S. Bojer, Valeria Bortolaia, Martine C. Holst Sørensen, Lone Brøndsted

## Abstract

Novel solutions are needed to reduce the risk of transmission of extended spectrum β-lactamase and AmpC β-lactamase producing *Escherichia coli* (ESBL/AmpC *E. coli*) in livestock to humans. Since phages are promising biocontrol agents, we established a collection of 28 phages against ESBL/AmpC *E. coli* and showed by whole genome sequencing that all phages were unique and could be assigned to 15 different genera. Host range analysis showed that 82% of 198 strains, representing the genetic diversity of ESBL/AmpC *E. coli*, were sensitive to at least one phage. Identifying receptors used for initial binding experimentally as well as *in silico* predictions, allowed us to combine phages into two different cocktails with broad host range targeting diverse receptors. These phage cocktails inhibit growth and kill ESBL/AmpC *E. coli in vitro*, thus suggesting the potential of phages as promising biocontrol agents.

**HIGHLIGHTS:**

- 28 unique phages infecting ESBL/AmpC *E. coli* were isolated and characterized
- Broad host range phages targeting different receptors were used to compose phage cocktails
- Phage cocktails efficiently inhibit growth of ESBL/AmpC *E. coli in vitro*

## INTRODUCTION

The rise of the antibiotic resistance and emergence of multi-drug resistant bacteria is a global problem. Of special concern are extended spectrum β-lactamase (ESBL) and AmpC β-lactamase (AmpC) producing *E. coli* (hereafter referred to as ESBL/AmpC *E. coli*) that are resistant to a broad spectrum of antibiotics, including penicillin and third generation cephalosporins ^1^. ESBL/AmpC *E. coli* show large genomic diversity and are represented in all phylogroups and sequence types (STs) of *E. coli* ^2–4^. While the AmpC β-lactamases are chromosomal encoded genes with upregulated expression ^5^, ESBL genes are mainly associated with a wide range of endemic conjugative plasmids ^6–9^. Due to the conjugative nature of these plasmids, ESBL genes may spread both *in vitro* and *in vivo*^10^. In livestock ESBL/AmpC *E. coli* is mainly commensal but may transfer antibiotic resistance genes into pathogenic *E. coli* as well as related pathogens to the human reservoir through contaminated foods ^11,12^. Thus, applying a One Health approach reducing the prevalence of ESBL/AmpC *E. coli* in animal reservoirs may minimize emergence of antibiotic resistant pathogenic *E. coli* ^13^. Different decolonization approaches have been proposed to reduce ESBL/AmpC *E. coli* prevalence in poultry flocks and pig pens ^14^. Among these approaches are diverse cleaning and disinfection agents, attempting competitive exclusion using probiotic cultures, and specific feed additives showed strong effect in prevention in some studies (reviewed in ^15^. However, complete decolonization of animals has proven challenging, and, in most cases, the applied approaches were ineffective against ESBL/AmpC *E. coli* ^14,16^. There is therefore a need for alternative methods to reduce the numbers of ESBL/AmpC *E. coli* in livestock.

Bacteriophages (phages) are viruses that infect and kill bacteria and have been used for biocontrol purposes as well as phage therapy targeting pathogenic bacteria (reviewed in^17^). Phages are host-specific, often infecting only specific species or even strains, leaving the rest of the microbiota unharmed. Additionally, they are self-replicating and self-limiting as they replicate only in the presence of a suitable host ^18^. A large number of diverse phages infecting *E. coli* have been described and many collections are well characterized, providing an insight into their diversity, genomics, classification, and interactions with their *E. coli* host ^19–26^. These studies show that coliphages are found in many different environments, including faeces, wastewater, soil, and water and have diverse host ranges infecting specific *E. coli* strains. Still, only a few studies using existing coliphage collections with limited characterization have been used to determine phage susceptibility of ESBL/AmpC *E. coli*. Two studies showed that phages isolated using environmental *E. coli* strains can infect and kill ESBL/AmpC *E. coli* with varying but low efficiency ^27,28^. In addition, we previously determined the ability of 16 coliphages isolated on the *E. coli* Reference collection (ECOR) to infect and form plaques on a large diverse ESBL/AmpC *E. coli* collection ^4^. However, these phages were only able to infect 23% of the 198 strains in the collection with varying efficacy, thus suggesting a need for other phages to cover the diversity of ESBL/AmpC *E. coli* ^4^. Other studies isolated phages using ESBL/AmpC *E. coli* as hosts, but only determined lysis on bacterial lawns adding phages in high concentration and true phage infection were not demonstrated ^29–32^. Thus, studies characterizing phage infection of ESBL/AmpC *E. coli* as well as their potential for biocontrol are limited. To increase the diversity of phages infecting ESBL/AmpC *E. coli* and provide well-characterized phages for biocontrol, we isolated and characterize phages infecting diverse ESBL/AmpC *E. coli* and used our collection to compose two different phage cocktails to explore the possibility of using them for biocontrol of ESBL/AmpC *E. coli*.

## RESULTS

### Isolation of phages infecting ESBL/AmpC *E. coli* using a diverse set of strains and samples

For isolating phages infecting ESBL/AmpC *E. coli*, we took advantage of our previously established large collection of 198 ESBL/AmpC *E. coli* covering the genetic diversity of this group of bacteria ^4^. To maximize the chances of isolating diverse phages, we selected 19 diverse strains as isolation hosts based on their different genetic background defined by multi-locus sequence typing (MLST) and carriers of diverse β-lactamase genes and plasmids (Figure 1). The samples for phage isolation were collected from environments expected to contain ESBL/AmpC *E. coli* including five samples of pig waste from a biogas production plant and two samples of broiler faeces. Additionally, two samples from aeration tanks in a wastewater treatment plant were collected to increase the likelihood of capturing diverse phages, as wastewater has proven to be a rich source of phages ^24^. A total of 28 phages were isolated either by direct plating or by enrichment in the presence of one of the 19 different isolation host (Figure 1). Most phages were isolated from wastewater (n=14) and pig waste (n=12), whereas only two phages were isolated from broiler faeces. There were no apparent correlations between the origin of the isolation hosts and the phages isolated from samples, as host strains originating from pig mostly isolated phages from wastewater and two phages from pig waste, whereas broiler strains isolated phages from all sources. Finally, most strains isolated only 1-2 phages, thus indicating potentially diverse phages.

**Figure 1:**
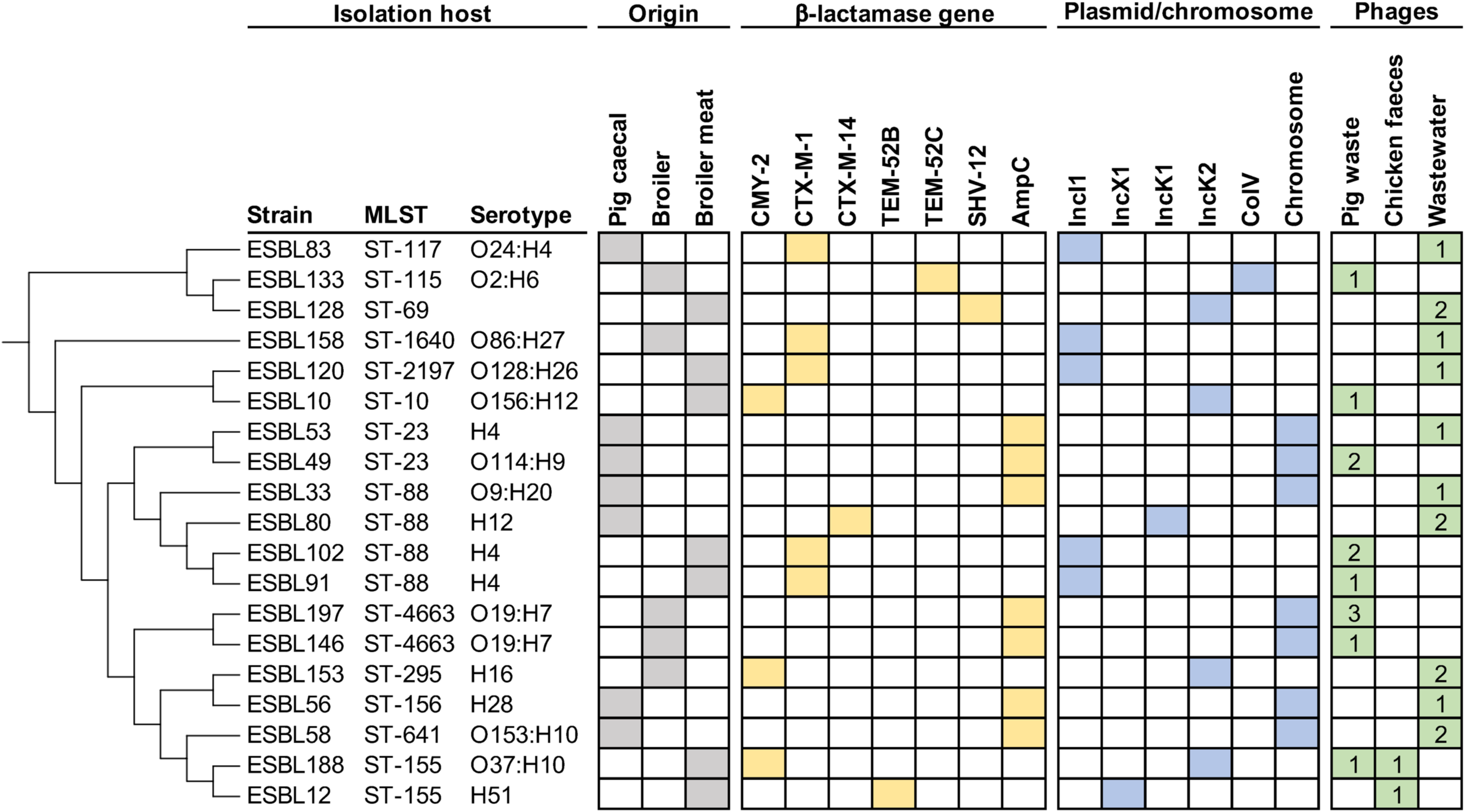
Diverse ESBL/AmpC *E. coli* are used for phage isolation from animal and wastewater samples. The MLST was extraxted from Enterobase and a phylogenetic tree was made and visualized using iTOL v6 software (Letunic and Bork, 2021). Columns indicate ST using MLST, serotypes, origin of strain (grey), type of β-lactamase gene (yellow) carried on specific plasmid types or the chromosome (blue) obtained from Vitt et al. 2023. Number of phages isolated from either pig waste, chicken faeces or wastewater on each isolation host is indicted (green).

### Taxonomic assignment and overall genetic comparison of ESBL/AmpC *E. coli* phages

To assign the phages into current taxonomic phage genera, all 28 isolated phages were whole genome sequenced and compared to existing phage genomes at the NCBI database using whole genome BLAST similarity search. Our analyses showed that the phages could be assigned to five different families: *Ackermannviridae*, *Autographiviridae*, *Drexlerviridae* and *Straboviridae* as well as several different subfamilies and genera. Among those, *Straboviridae* were the most abundant family, represented by 16 phages belonging to the genera *Krischvirus*, *Mosigvirus* and *Tequatrovirus*. The majority of the 28 phages showed high nucleotide similarity (96-99% identity, 92-98% coverage) to other phages infecting *E. coli* (Table 1). Exceptions were *Rosemountvirus* AV127 showing similarities to phages infecting *Salmonella* and phage AV124 to *Klebsiella* phage Seu621 belonging to the genus *Mydovirus*. The lowest sequence similarity to other known phages was observed for phage AV104, sharing only 88% nucleotide identity over 75% coverage to its closest relative phage fFiEco02, a *Kagunavirus* infecting *E. coli*. Thus, while a few phages were genetically distinct from previously sequenced phages, most of the phages in our collection are genetically related to known phages.

**Table 1:**
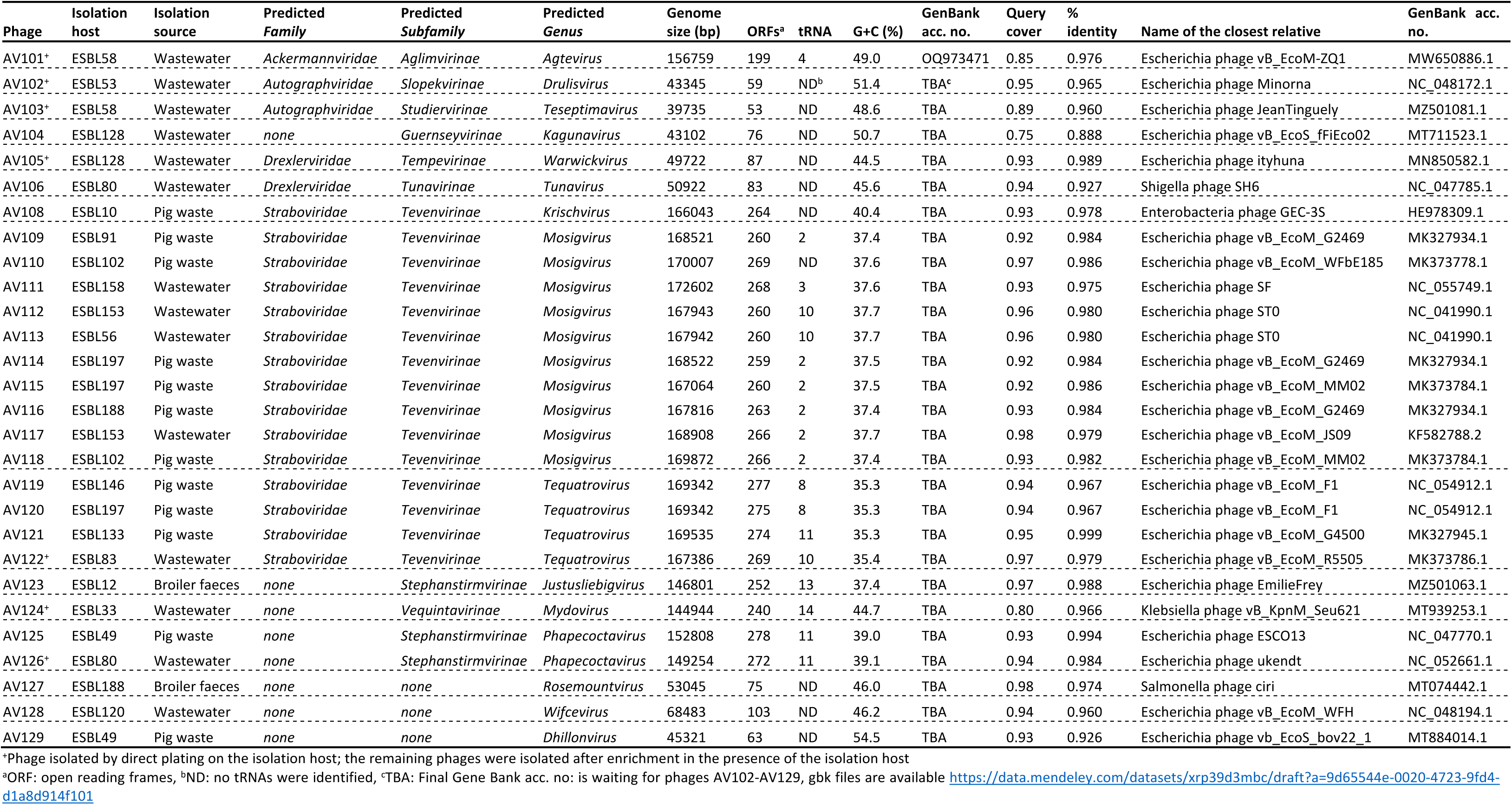
Genome characteristics of phages infecting ESBL/AmpC *E. coli*.

To understand the genetic diversity within the phage collection, we compared all genomes at nucleotide level constructing an identity matrix based on the WGS data (Figure 2). This analysis showed that the phages are distinct from each other and confirmed that they cluster according to their predicted taxonomic classification. Further genomic analysis revealed varying number of ORFs, tRNAs as well as GC content associated with the assigned genus (Table 1). Functional annotation of the larger group of phages belonging to the subfamily of *Tevenvirinae* demonstrated high overall similarity and synteny in genome organization (Figure 3A and 3B). These phages contain the typical features of *Tevenvirinae* including genome sizes of 162-250 kb, a genomic organization of clusters of early, middle, and late genes, a varying number of homing endonucleases, the presence of several tRNAs and hyper modification of cytosine residues to protect the phage against different restriction-modification systems of the host ^33–35^. Comparative genomics of *Tevenvirinae* phages within *Mosigvirus* and *Tequatrovirus* showed minor variations in all parts of the genome. In addition, the region encoding the long tail fiber gp37 and the gp38 adhesins known recognize host receptors in phages T4 and T2 and T6, respectively varies between phages (Figure 3B and 3C). Finally, functional annotation of the more rarely isolated *Phapecoctavirus* phages AV125 and AV126 identified large numbers of uncharacterized ORFS and hypothetical genes (253 out of 278 genes are annotated as hypothetical genes in AV125) as well as only few differences between these two phages including a putative receptor binding protein (Figure 3D). None of the phages in the collection encode integrases or potential repressors to maintain lysogeny, suggesting that all phages are virulent. Finally, screening all phage genomes for virulence factors and antibiotic resistance genes using program VirulenceFinder ^36^ suggested that none of the phages encode virulence genes and thus may be for biocontrol applications targeting ESBL/AmpC *E. coli*.

**Figure 2:**
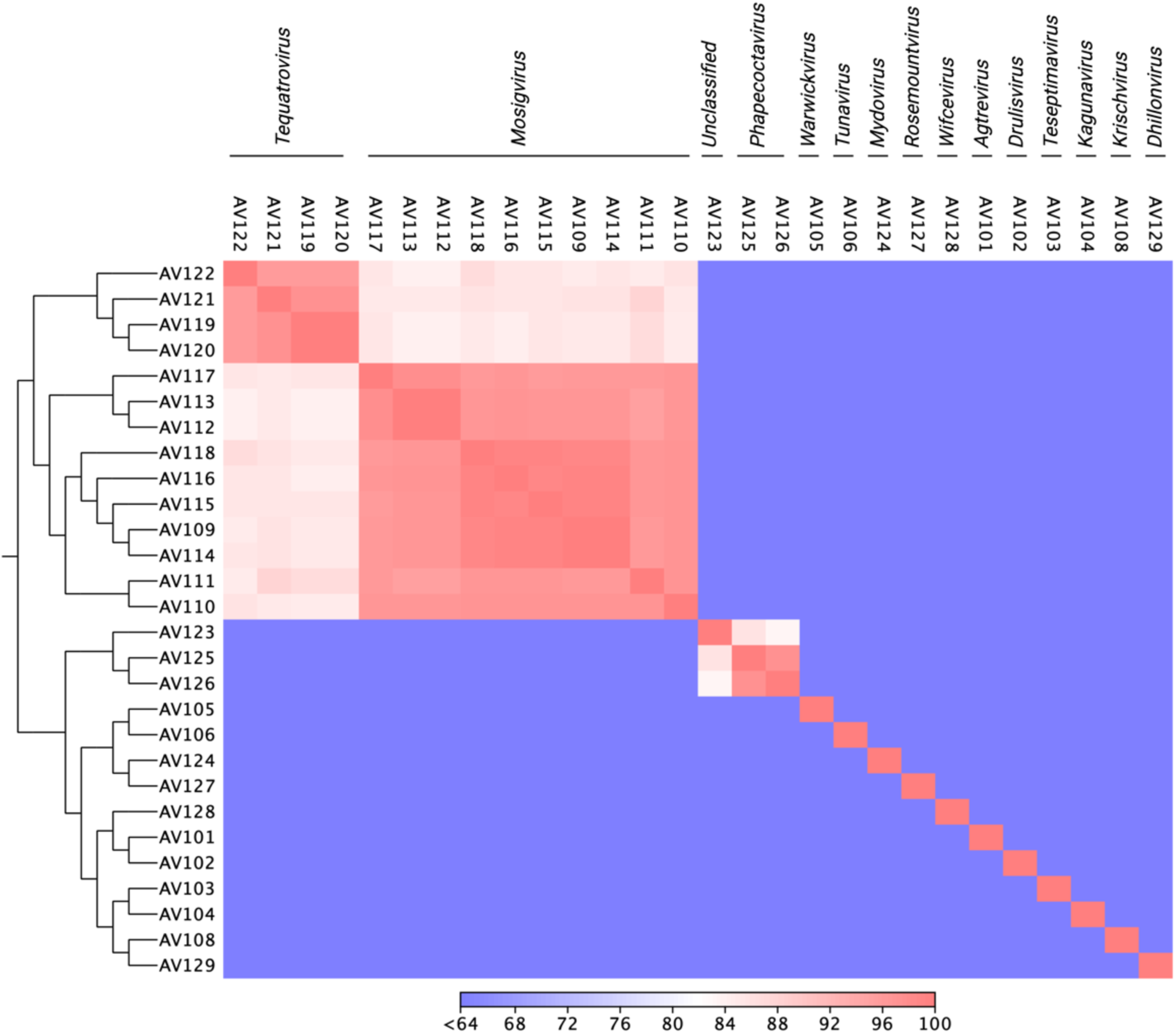
Genomic similarity among isolated phages infecting ESBL/AmpC *E. coli.* Alle phage genomes were aligned, and the average nucleotide identity (ANI) was calculated for each pair of genomes based on all aligned regions of the whole genome alignment. The colour bar indicates ANI as the percentage of exactly matching nucleotides.

**Figure 3:**
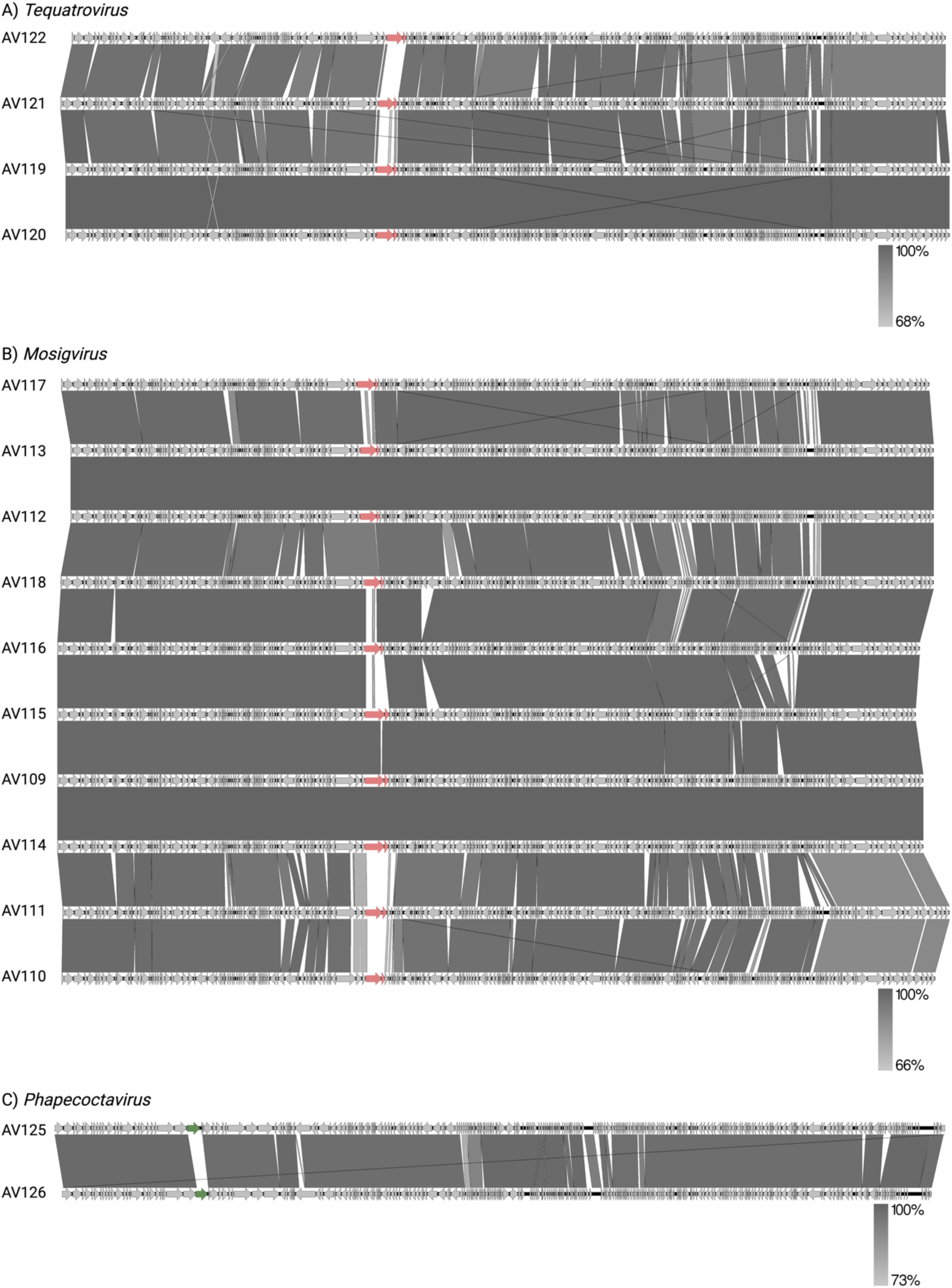
Comparative genomics of phages infecting ESBL/AmpC *E. coli* belonging to (A) *Mosigvirus*, (B) *Tequatrovirus* and (C) *Phapecoctavirus*. Putative receptor binding proteins as *gp37* and *gp38* for Mosigvirus and *Tequatrovirus* as well as diverse genes in the tail fibre locus of *Phapecoctavirus* are indicated by red arrows.

### Phage host range of a large collection representing the diversity of ESBL/AmpC *E. coli*

To determine the ability of the isolated phages to infect diverse ESBL/AmpC *E. coli*, we performed an extended host range analysis determining plaque formation using our large collection of ESBL/AmpC *E. coli*. This collection contains 198 commensal ESBL/AmpC *E. coli* from two animal reservoirs including pigs and broilers and the meats hereof, representing all known phylogroups as well as 65 sequence types (STs) (Vitt et al., 2023). Despite the genomic diversity of the ESBL/AmpC *E. coli* collection, a total of 162 strains (82 % of all 198 strains) showed susceptibility to at least one phage in our collection (Figure 4). Among the sensitive ESBL/AmpC *E. coli* were strains belonging to phylogroups A, B1 and C that are mainly represented by commensal *E. coli* but importantly also strains within the phylogroups B2, D, E, F, G, and clade I that usually carry virulence genes and are potentially pathogenic ^37^ (Figure 4). For example, *Mosigvirus* were the only phages infecting strains of phylogroup G, while strains in phylogroup B2 were only infected by *Tequatrovirus*, *Justusliebigvirus* and *Phapecoctavirus* phages (Table S2). Overall, the host range of the entire phage collection span across most STs and phylogroups thus demonstrating phage infection of genetically diverse ESBL/AmpC *E. coli*.

**Figure 4.**
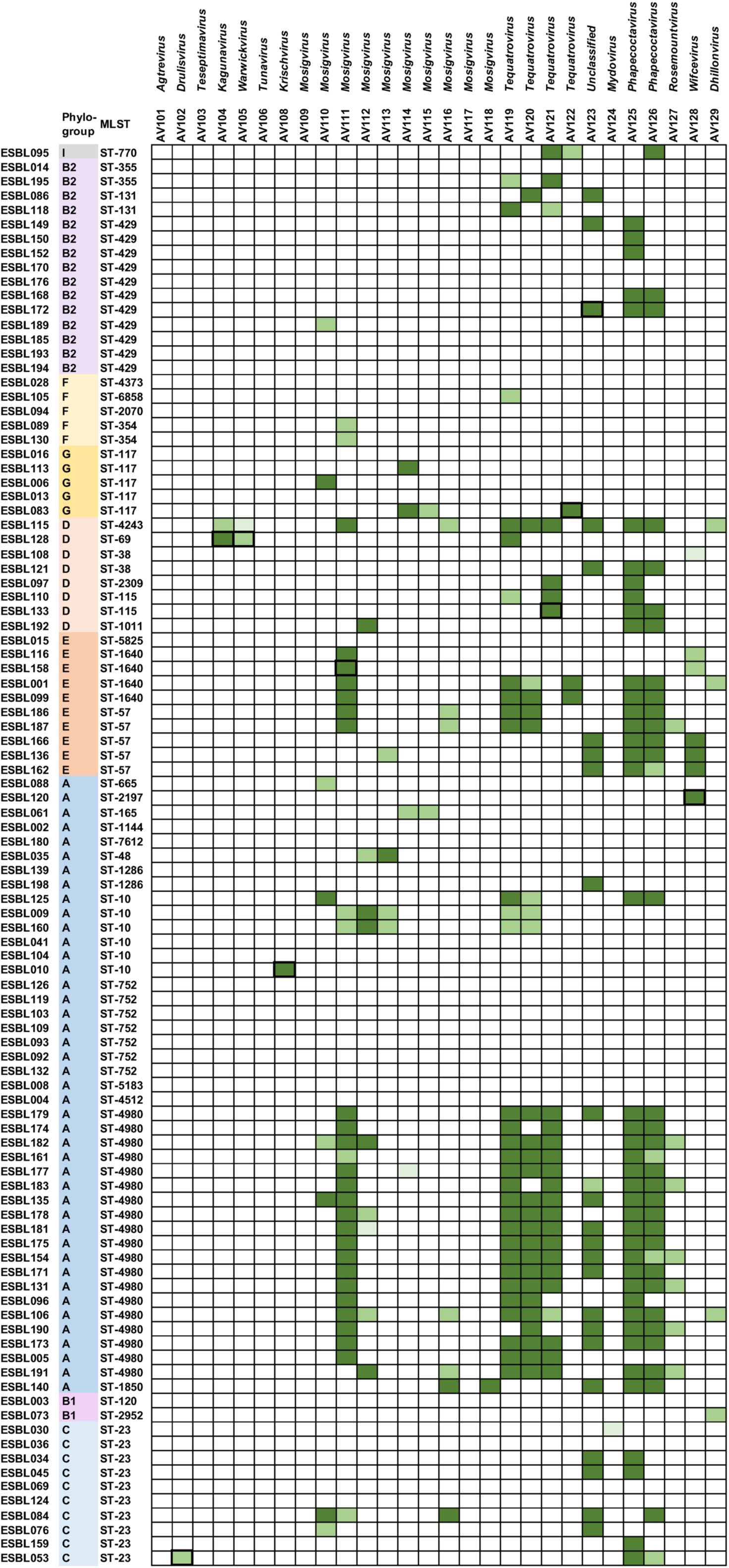
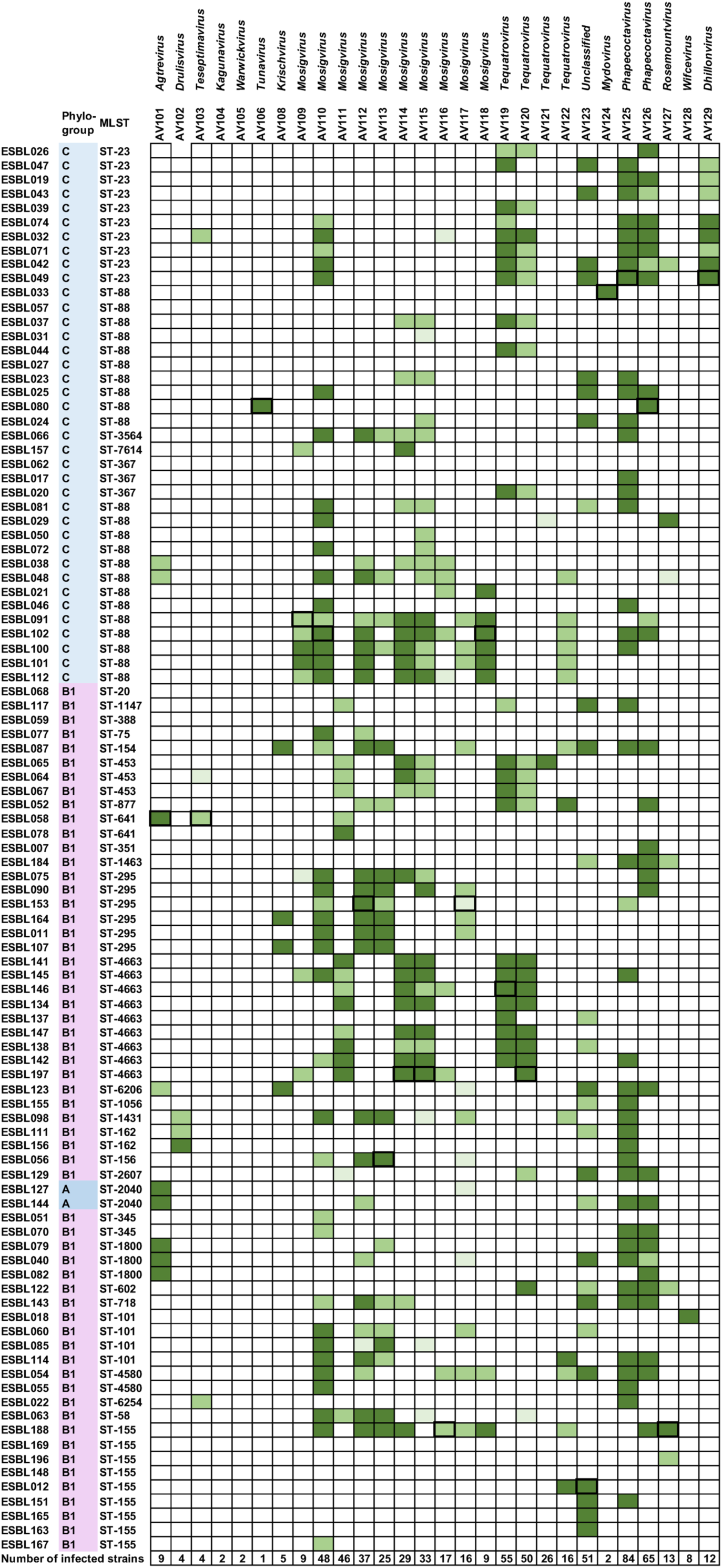
Phage host range analysis using plaque assay on 198 diverse ESBL/AmpC *E. coli*. Strains are grouped according to phylogeny and columns indicate phylogroups marked with different colours and STs using MLST based to our previous analysis (Vitt et al. 2023). Isolation hosts are indicated by a black box. More than 10^8^ pfu per ml (dark green). Between10^5^ to 9.99×10^7^ pfu per ml (medium green). Less than 9.99×10^4^ pfu per ml (light green). No plaques observed (white).

The host range analysis also demonstrated that each phage showing unique host range profile. While some phages were infecting only one or a few ESBL/AmpC *E. coli*, others formed plaques on nearly 40 % of the entire strain collection (Figure 4). Several phages like for example *Agtrevirus* AV101, *Mosigvirus* AV109 and *Wificvirus* AV127 showed narrow host ranges with specificity only towards their isolation host and a few other strains (Figure 4). In contrast, other phages showed broad host ranges infecting ESBL/AmpC *E. coli* across most phylogroups and many different STs (Figure 4; Table S2). For example, *Mosigvirus* AV110 and AV111, *Tequatrovirus* AV119 and AV120 as well as *Justusliebigvirus* AV123 infects between 46 to 55 strains of ESBL/AmpC *E. coli* (Figure 4). Finally, *Phapecoctavirus* AV125 and AV126 displayed the broadest host range of all phages in the collection, infecting 84 and 65 ESBL/AmpC *E. coli*, respectively (Figure 4). Remarkably, these phages were not only infecting closely related strains but also a wide range of genetically diverse isolates belonging to 28 to 35 different STs and six phylogroups (Table S2). Overall, the broad complementary and diverse host ranges suggest that single phages may be combined into a cocktail covering the majority of ESBL/AmpC *E. coli* diversity.

### Receptor identification

To determine the bacterial receptors used for initial phage binding, we spotted ten-fold serial dilutions of phage stocks on lawns of wild type *E. coli* ECOR4 and as well as defined deletion mutants of known phage receptors (Figure 5A). This identified the phage receptors among outer membrane proteins BtuB, OmpA, OmpC, and Tsx as well as lipopolysaccharide (LPS) for nine phages that were able to infect wildtype *E. coli* ECOR4 (Figure 5A). To propose receptors for the remaining *Tevenvirinae* phages, we extracted sequences of predicted receptor binding proteins (RBPs) and performed BLASTp analysis. The well-studied *Tevenvirinae* phage T4 uses long tail fibers (Gp37) to interact with the host receptor^38^. In addition, in phage T4 Gp38 serves as chaperone for Gp37 folding, whereas Gp38 homologues of related *Tevenvirinae* phages T2 and T6 encode an adhesin responsible for binding to the bacterial receptors ^39^.Using amino acid alignment, we demonstrated that seven phages encode a chaperone showing 83-100% similarity to Gp38 of T4, suggesting use of Gp37 long tail fibers for host recognition (Figure S1). The remaining *Tevenvirinae* eight phages showed similarity to the Gp38 adhesin of T2 and T6 responsible for host recognition (Figure S2).

**Figure 5:**
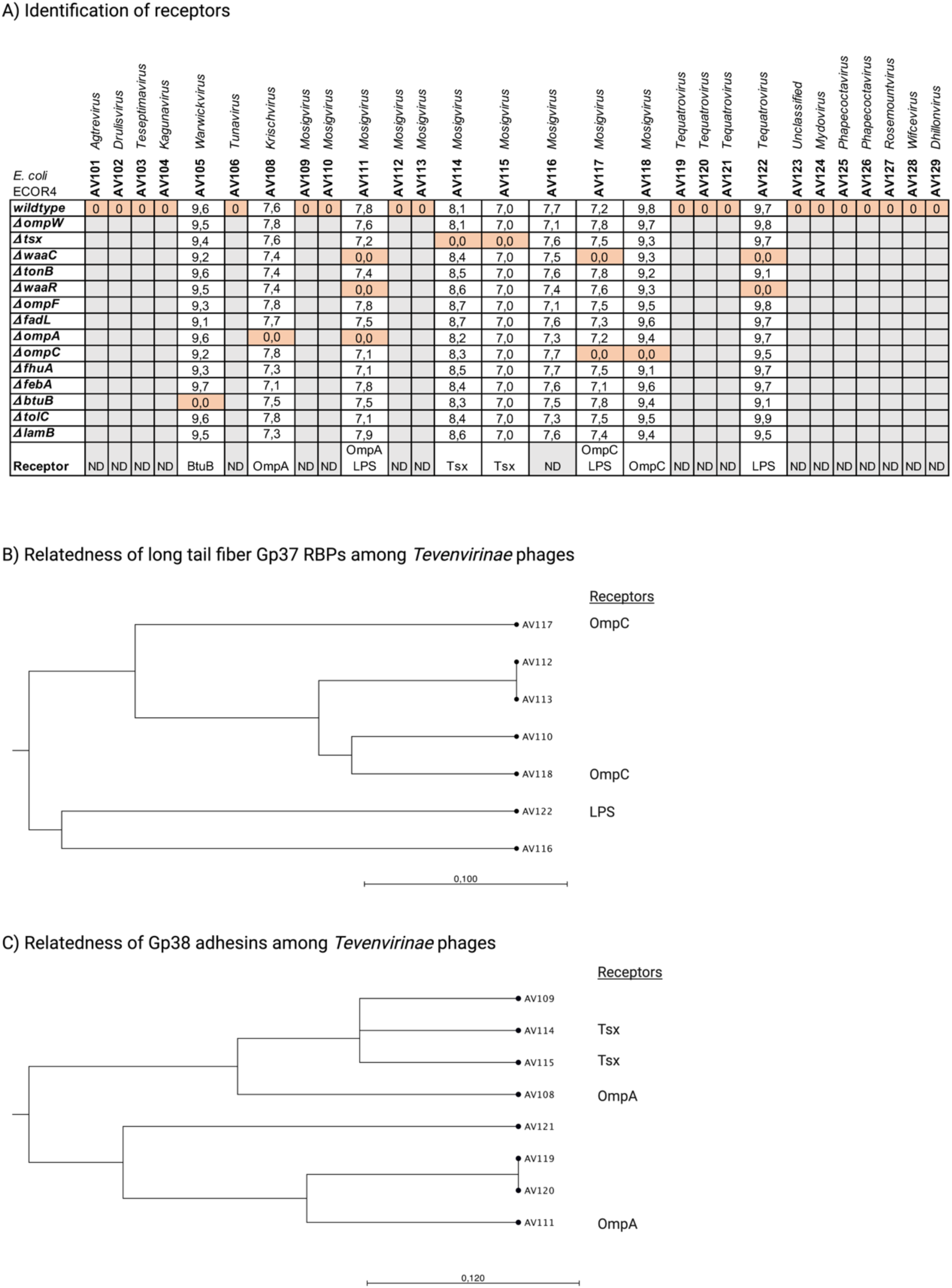
Identification of bacterial receptors. (A) Log10(PFU per ml) obtained from plaque assay on lawns of *E. coli* wildtype ECOR4 as well as ECOR4 deletion mutants as indicated. (B) Phylogenetic relatedness of long tail fibers gp37 of *Mosigvirus* phages AV110, AV112, AV113, AV116, AV117 and AV118 as well as *Tequatrovirus* phage AV122. (C) Phylogenetic relatedness of gp38 adhesins of *Krischvirus* phage AV108, *Mosigvirus* phages AV109, AV111, AV114 and AV11 as well as *Tequatrovirus* phages AV119, AV120 and AV121.

*Mosigvirus* phages AV110, AV112, AV113, AV116, AV117 and AV118 as well as tequatrovirus AV122 encode a Gp38 chaperone, hence predicted to use their Gp37 long tail fibers to bind their receptors. Alignment analysis showed that the major differences of Gp37 were observed within the head domain (corresponding to 918-973 of gp37 of T4) of the tail tip, previously demonstrated to be responsible for receptor binding of phage T4 ^40^ (Figure S3). Phylogenetic analysis indicated that Gp37 of phages AV110, AV112, and AV113 may bind to OmpC, as confirmed experimentally for phages AV117 and AV118 (Figure 5A and B). In contrast, phages AV116 and AV122 may be dependent on LPS for infection, but an unidentified outer membrane receptor cannot be ruled out as a secondary receptor (Figure 5B).

The Gp38 adhesin of *Tevenvirinae* phages T2 and T6 consist of an N-terminal attaching to the long tail fibers and a C-terminal comprised of five conserved glycine-rich motifs (GRMs) and four hypervariable segments (HVSs) ^39^. Alignment analysis demonstrated that mosigvirus AV111 as well as tequatrovirus AV119, AV120 and AV121 are most similar to the Gp38 adhesin of T6, whereas Krischvirus AV108 and mosigvirus AV109, AV114 and AV115 showed similarity to the Gp38 adhesin of T2 (Figure S2). In general, the main differences among all Gp38 homologues were found within the HVSs responsible for host recognition (Figure S2). Phylogenetic analysis showed that the Gp38 adhesin of phages AV119, AV120 and AV121 are most closely related to Gp38 of phage AV111 dependent on OmpA for infection, suggesting that these phages also bind to OmpA (Figure 5A and 5C). Phylogenetic analysis confirmed that the Gp38 adhesin of phage AV109 is closely related to phages AV114 and AV115, shown to recognize Tsx, thus suggesting that phage AV109 may use this receptor as well (Figure 5A and 5C). In contrast, phage AV108 uses OmpA as receptor and while its Gp38 adhesin showed similarities to the adhesins of AV109, AV114 and AV115, differences within the HVS3 and HVS4 may be responsible for binding to two different receptors (Figure S2). Overall, our phage collection may target at least six different receptors, some predicted by bioinformatic analysis while others were demonstrated experimentally.

### Phage-mediated growth inhibition of ESBL/AmpC *E. coli in vitro*

To demonstrate the therapeutic potential of our phages, we tested the ability of selected individual phages to inhibit growth *in vitro* of two different strains; ESBL102 that carries a CTX-M-1 β-lactamase on an IncI1 plasmid and ESBL145 expressing a chromosomal upregulated AmpC. Moreover, ESBL102 and ESBL145 belongs to phylogroups C and B1, as well as ST-88 and ST4663, respectively, thus representing diverse ESBL/AmpC *E. coli*. At MOI of 10, we observed that single phages AV110, AV111, AV114, AV118 and AV125 may inhibit growth for up to eight hours whereafter growth was initiated probably due to development of resistance (Figure 6A and 6B). At lower MOIs similar patterns were observed, but with a tendency of an earlier onset of growth and thus resistance development (data not shown). Thus, to prevent resistance development, we designed two phage cocktails consisting of phages targeting different receptors. In both cases we included phapecoctavirus AV125 in the cocktail, as this phage infects the most strains (n=84) (Figure 3) in the collection. A remarkable feature of the *Stephanstirmvirinae* phages like phapecoctavirus AV125, is the presence of four different sets of tail fibers and two tail spike proteins that form a structure resembling an open “nanosized Swiss army knife” with tail fibers pointing in three directions ^41^. Such tail structure is suggested to provide broad host specificity proposed to target polysaccharides of the capsule, enterobacterial common antigen and lipopolysaccharides as receptors ^42^. We composed one cocktail of mosigvirus AV114 (infecting 29 strains using Tsx as receptor), mosigvirus AV118 (infecting 9 strains using OmpC as receptor) and phapecoctavirus AV125. The second cocktail was composed of mosigvirus AV110 (infecting 48 strains with the predicted receptor OmpC), mosigvirus AV111 (infecting 46 strains using OmpA as receptor), mosigvirus AV114 (infecting 29 strains using Tsx as receptor), and phapecoctavirus AV125. Interestingly, we found that both cocktails were able to inhibit growth of their target strain for entire 24 hours of the experiment (Figure 6C and 6D). Overall, the results suggested that phage cocktails targeting different receptors can be used to inhibit the growth of ESBL/AmpC *E. coli in vitro* and could potentially be used for biocontrol.

**Figure 6:**
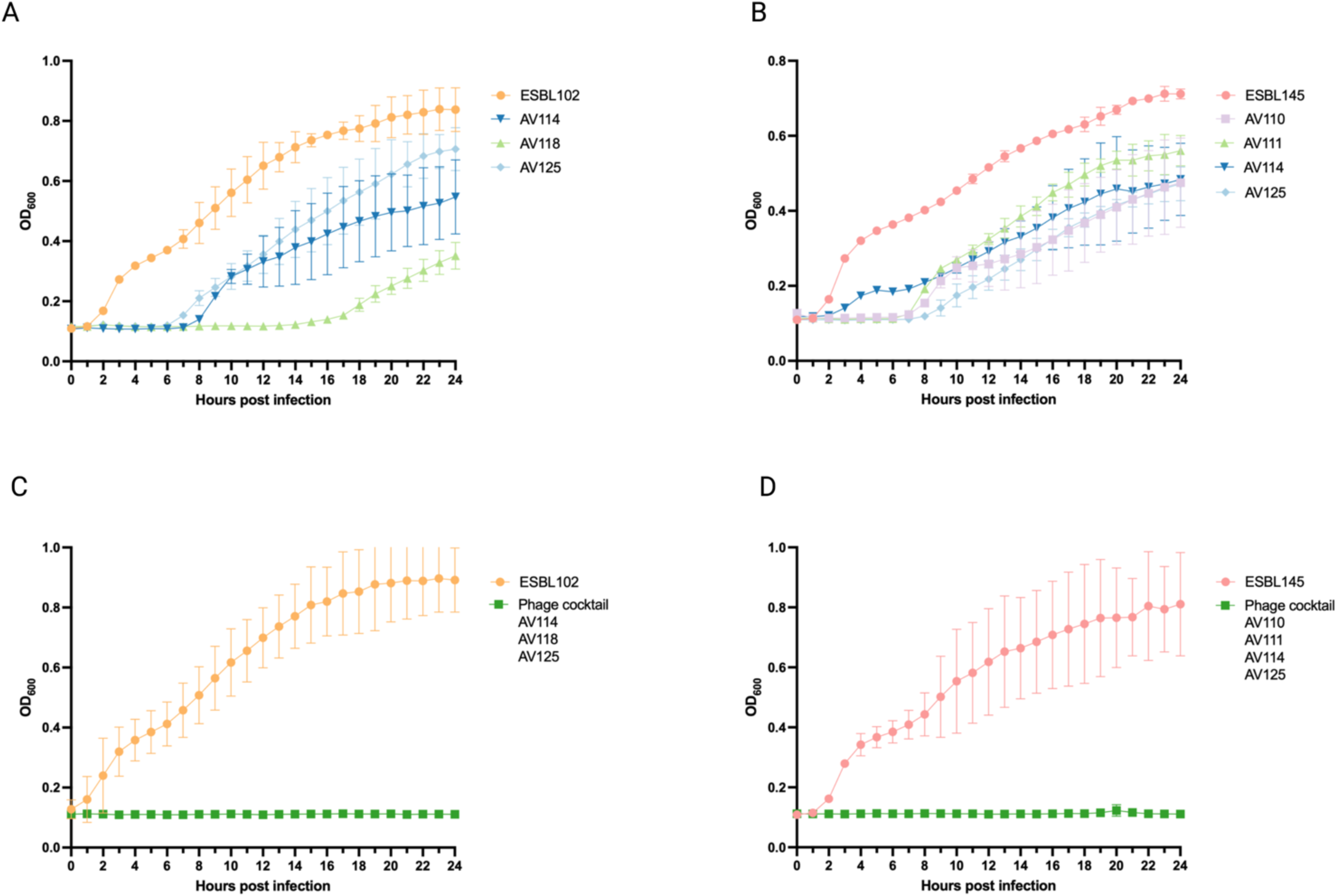
Phage-mediated growth inhibition of ESBL/AmpC *E. coli in vitro.* (A) Growth of ESBL102 in the absence (orange) of phages and presence of phage mosigvirus AV114 (dark blue), mosigvirus AV118 (light green) and phapecoctavirus AV125 (light blue). (B) Growth of ESBL145 in the absence (light red) of phages or in the presence of mosigvirus AV110 (light purple), mosigvirus AV111 (light green), mosigvirus AV114 (dark blue), and phapecoctavirus AV125 (light blue). (C) Growth of ESBL102 in the presence (dark green) or absence (orange) of phage cocktail consisting of mosigvirus AV114 (Tsx receptor), mosigvirus AV118 (OmpC receptor) and phapecoctavirus AV125. (B) Growth of ESBL145 in the presence (dark green) or absence (light red) of phage cocktail consisting of mosigvirus AV110 (predicted OmpC), mosigvirus AV111 (OmpA receptor), mosigvirus AV114 (Tsx receptor), and phapecoctavirus AV125.

## DISCUSSION

The rise of antibiotic resistance is a global problem within human as well as veterinary medicine and resistance to broad spectrum β-lactam antibiotics is found among highly diverse ESBL/AmpC *E. coli*. As livestock are one of the main reservoirs of ESBL/AmpC *E. coli* there is a risk of transmission of resistant bacteria as well as antibiotic resistance genes to farmers and consumers through direct contact to animals and food. Taking a One Health approach reducing ESBL/AmpC *E. coli* in kivestock and foods will positively impact animal and human health as well as the environment. As phages may be a promising approach to reduce ESBL/AmpC *E. coli*, we established and characterized a collection of 28 diverse phages targeting a broad range of ESBL/AmpC *E. coli* representing the genetic diversity found worldwide ^4^ and demonstrate application of phages as a promising approach for biocontrol of ESBL/AmpC *E. coli*.

Although most of our phages fall into established genera of coliphages, exhibiting high overall nucleotide similarity over 90%, they are yet distinct from their closest relatives and genus/subfamily representatives, thus expanding on coliphage diversity. In addition, the closest relatives of several phages showed similarities to other phages isolated from wastewater ^22,24,25^, highlighting the abundance of these phages in wastewater treatment plants across different geographical locations. Interestingly, two phages showed low overall nucleotide sequence similarity to known phages. These were AV124 and AV104 classified within the *Vequintavirinae* subfamily (likely *Mydovirus* genus) and *Kagunavirus* of *Drexlerviridae*, respectively, that showed low overall nucleotide similarity of 77% (96.5% identity, 80% coverage) and 66% (88.8% identity, 75% coverage) to their closest relatives. Interestingly, the few known kagunavirus were identified through metagenomics data originating from microbiome studies ^43^. Phage AV124, on the other hand, only had close relatives of phages infecting *Klebsiella*, thus likely explaining its narrow host range within the ESBL/AmpC *E. coli*. Similarly, another narrow host range phage AV127, belonging to *Rosemountvirus* genus, and showed similarity to *Salmonella* phages only. The hosts of the coliphages in our collection may therefore extend to other species of *Enterobactericeae* such as *Enterobacteria, Klebsiella, Salmonella* and *Shigella,* based on the similarities to phages infecting these species, yet most of the closely related phages infect *E. coli*.

Phages for biocontrol or phage therapy must be specific against the target bacterium and preferably have a broad host range to cover the diversity of the target bacterial population. Importantly, we showed that phages of our collection combined infects all phylogroups and most of the 65 ST groups of our diverse ESBL/AmpC *E. coli* collection, thus showing a broad coverage of the diversity of ESBL/AmpC *E. coli*. Furthermore, isolating phages specifically targeting ESBL/AmpC *E. coli* significantly increased the coverage from 23% in our previous study to 82% in our present work ^4^. Still, the overall susceptibility of ESBL/AmpC *E. coli* could be further improved by isolating phages using ESBL/AmpC *E. coli* know to be resistant to phage infection. Here we found that several phages within the *Tevenvirinae* subfamily as well as phages belonging to *Stephanstrimvirinae* had broad host ranges covering all phylogroups and various STs. To some extent, such wide host range may be attributed to protection against anti-phage defense mechanisms by genomic modifications, such as hyper modification of cytosine characteristic for *Tevenvirinae* phages also identified in our phages and rhamnose modification in *Stephanstrimvirinae* phages ^22,24,25,44^. While the cytosine hyper modification has proved to be efficient against RM-systems for phages of *Tevenvirinae* subfamily, some *Stephastirmvirinae* phages were shown to be sensitive to many RM-systems ^22,24,25,44^. Thus, suggesting that other mechanisms potentially encoded by some of the many unknown genes may allow successful infection of a wide host range of strains.

Phages are equipped with tail fibers or tail spikes that may allow recognition of diverse surface structures displayed by ESBL/AmpC *E. coli* ^45,46^. While *Stephanstirmvirinae* phages encode three conserved tail fibers carrying glycosidase and colanidase activity, some phages encode an additional tail spike with N-acetylneuraminidase activity but is not found in our phages (data not shown). So far, the specific receptors were not determined for our *Stephanstrimvirinae,* but other studies have suggested that these phages may initially bind to polysaccharides of enterobacterial common antigen and then to the outer core of lipopolysaccharide for DNA injection, however the role of the specific tail fibers in host binding have not been elucidated yet ^42^. The *Tevenvirinae* phages of our collection were encoding different tail fibers with similarity to homologues of either gp37 of T4 or the gp38 adhesin homologues of T2 or T6. The diversity of the tail fibers was mainly due to the variations within the hypervariable segments (HVSs) of the gp38 homologues of T2 or T6 and in the head domain of the tail tip of the homologues of gp37 of T4, known to influence phage binding to the host. In accordance, phages AV110, AV112, and AV113 encoding highly similar long tail fibers showed similar host range profiles, whereas the remaining phages using gp37 to bind their host showed differences in tail fibers as well as host ranges. Interestingly, phages AV111, AV119, and AV120 showed the broadest host range within our *Tevenvirinae* phages and were encoding highly similar gp38 adhesins possibly targeting OmpA. The host ranges were quite similar as well, but minor differences were observed that may be due to amino acid substitutions in HVS1 and the C-terminal of the three phages. Overall, the broad host range of some phages, like the *Justusliebigvirus, Phapecoctavirus* and *Tevenvirinae* phages suggest that they may be promising candidates for biocontrol of ESBL/AmpC *E. coli*.

Notably, a few studies have tested the efficacy of commercially available phage cocktails against ESBL/AmpC *E. coli*, including the Intesti bacteriophage cocktail consisting of at least 23 phages infecting different bacterial species causing intestinal, urinary tract and oral cavity infections caused by *E. coli* among other bacteria ^47^. Yet, the cocktail has not been tested systematically against ESBL/AmpC *E. coli.* Here, we designed two different phage cocktails and tested their ability to inhibit growth of ESBL/AmpC *E. coli in vitro* using knowledge of host ranges to select phages infecting the target strain. Similarly, for specific applications, phages for cocktails may be selected based on knowledge of the target ESBL/AmpC *E. coli* strain. To compose the most efficient phage cocktails, we used the obtained data to select phages that binds to diverse receptors when infecting ESBL/AmpC *E. coli*, thus reducing the chances of phage resistance development. Indeed, *in vitro* experiments demonstrated that the phage cocktails could inhibit the growth of ESBL/AmpC *E. coli* strains ESBL102 and ESBL145 over 24 hours without phage resistant development. In contrast, treatment by single phages lead to resistance development within up to 8 hours, thus demonstrating the power of phage cocktails in preventing resistance development ^48–52^. In conclusion, the present work demonstrates that phages in our collection are promising to target diverse ESBL/AmpC *E. coli* and have thus laid the foundation for further development for phage cocktails used for biocontrol of ESBL/AmpC *E. coli*.

### Limitations of the study

Here we describe isolation of 28 phages infecting ESBL/AmpC *E. coli* and provide a comprehensive host range analysis as well as receptor identification. However, further studies are needed to better understand the biology and potential applications of these phages, for example identification of the receptors of broad host range *Phapecoctavirus* as well as functional assignment of their unknown ORF. Future studies could as well focus on the host range determinants of phages in the collection, including if they infect other commensals *E. coli* that may be beneficial for the gut health or alternatively pathogenic strains of *E. coli*. Additionally, the potential for biocontrol could be further investigated by determining the ability of the phage cocktails to decolonize animals using mice models or farm animals as well as phage resistant development *in vivo*.

## STAR METHODS

Detailed methods are provided in the online version of the paper and include the following:

• KEY RESOURCE TABLE
• RESOURCE AVAILABILITY
  ◦ Lead contact
  ◦ Materials availability
  ◦ Data and code availability
• EXPERIMENTAL MODEL AND SUBJECT DETAILS
  ◦ Bacterial strains
• METHODS DETAILS
  ◦ Processing of animal and wastewater samples
  ◦ Phage isolation, purification, and propagation
  ◦ Phage plaque assay
  ◦ Host range analysis
  ◦ Determination of receptors in *E. coli* strain ECOR4
  ◦ Single bacteriophage and bacteriophage cocktail growth inhibition assay
  ◦ DNA extraction and sequencing
• QUANTIFICATION AND STATISTICAL ANALYSIS
  ◦ Genome assembly and analyses
  ◦ Bioinformatics analyses

## SUPPLEMENTARY INFORMATION

Supplementary information can be found online at (add link when available)

## ACKNOWLEDGEMENT

This project has received funding from Promilleafgiftsfonden, Denmark to ARV and ANS was supported by the Danish Council for Independent Research (2035-00112B).

## AUTHOR CONTRIBUTION

**Conceptualization**, A.R.V., M.C.H.S. V.B., A.N.S., and L.B; **Methodology**, A.R.V., A.N.S., and L.B; **Investigation**, A.R.V., A.N.S., and L.B.; **Resources**, L.B.; **Writing – Original Draft**, A.R.V. and L.B.; **Writing – Review & Editing**, A.N.S. and L.B; **Funding Acquisition**, L.B; **Resources**, L.B.; **Supervision**, M.C.H.S., V.B., and L.B.

## STAR METHODS

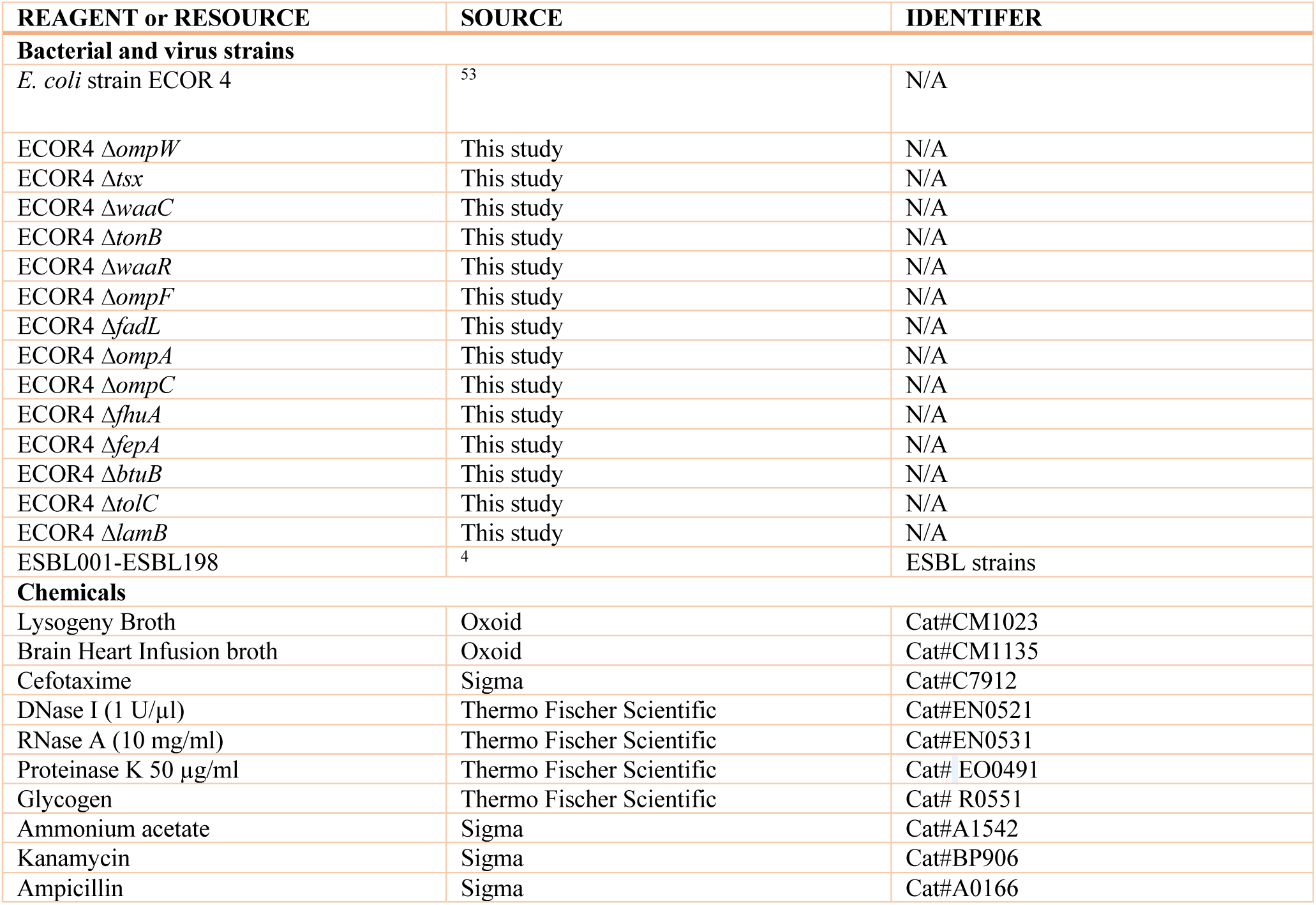
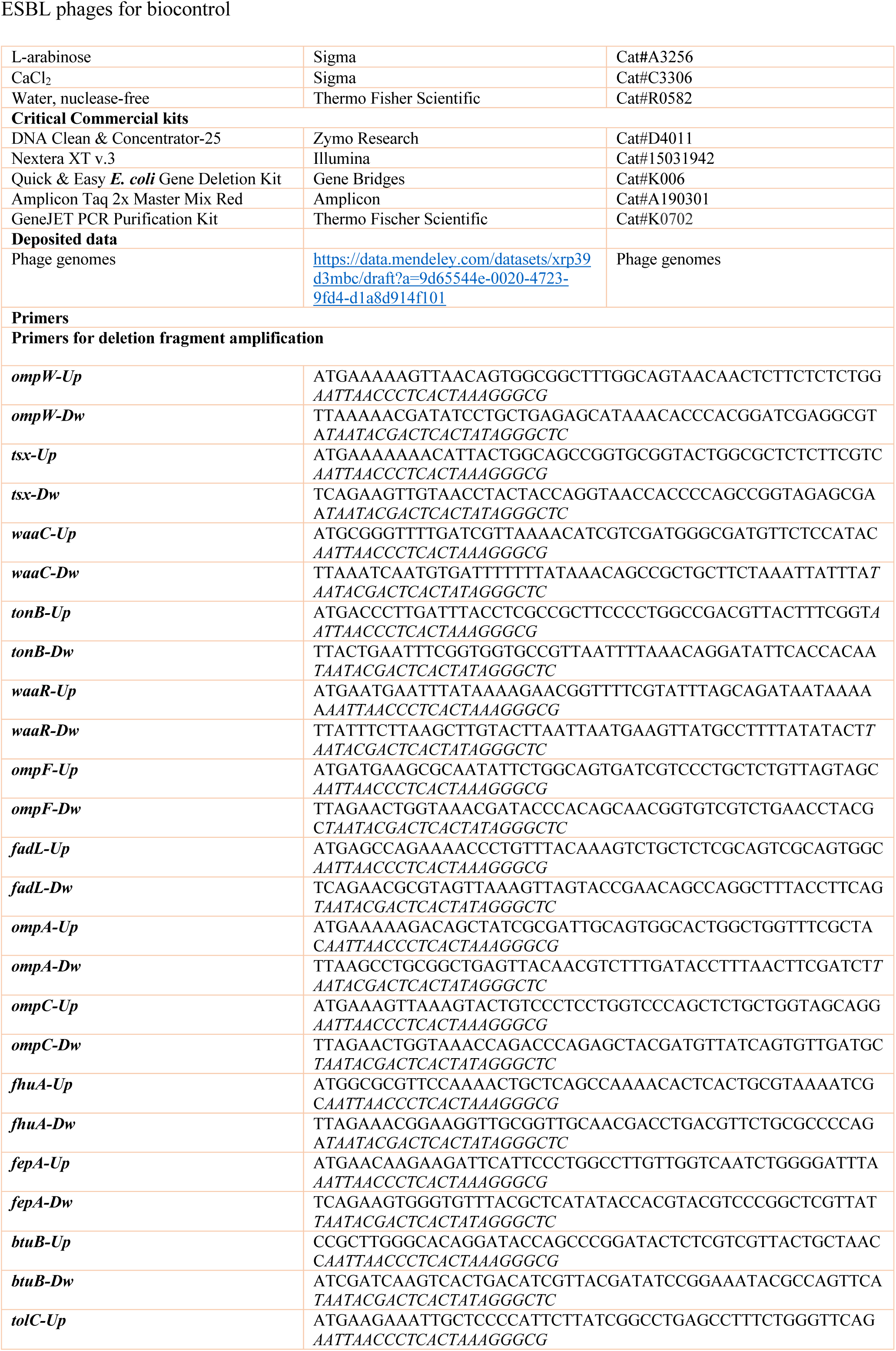
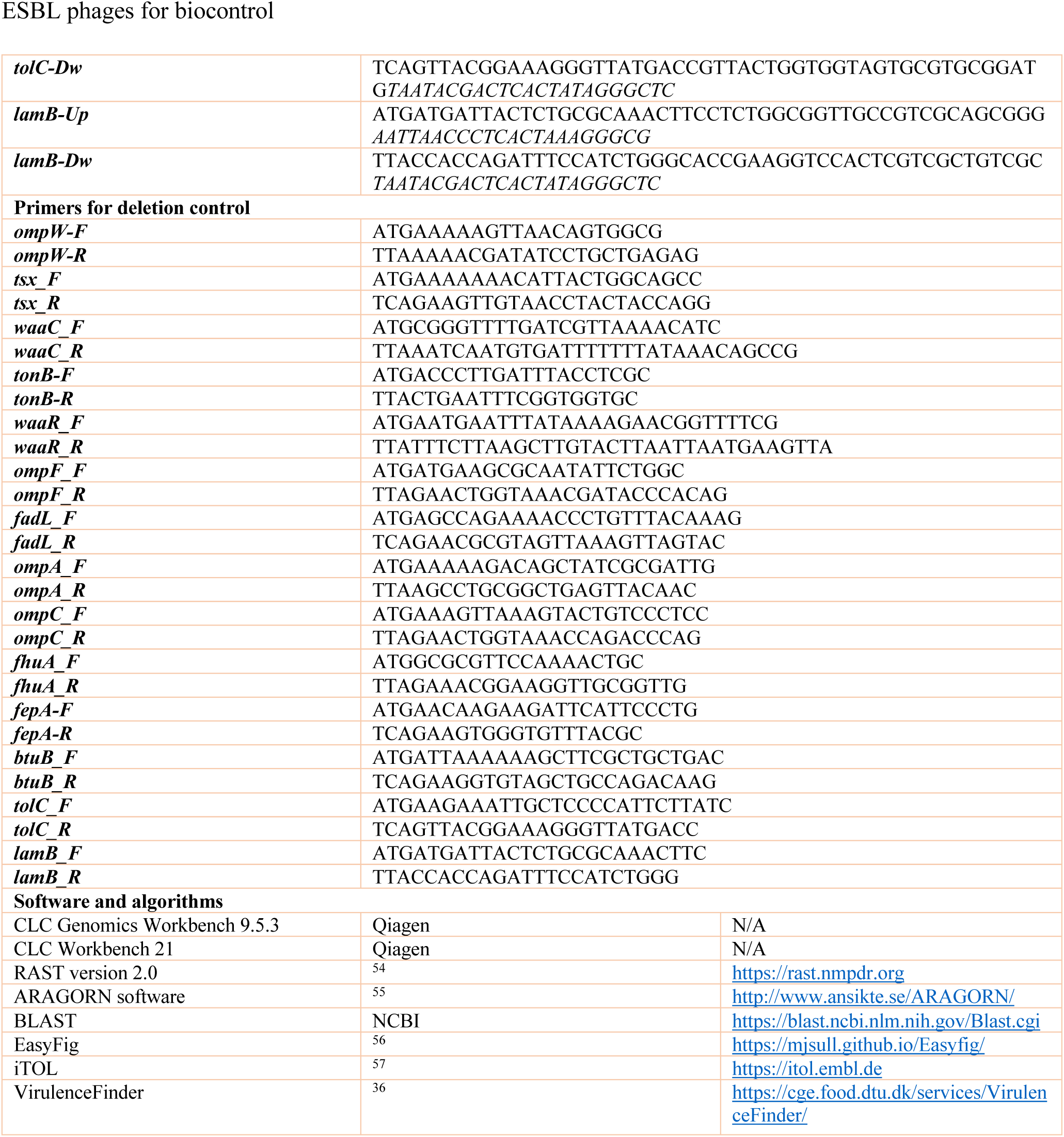
KEY RESOURCE TABLE.

### KEY RESOURCE TABLE

## RESOURCE AVAILABILITY

### Lead contact

Further information and requests for resources and reagents should be directed to and will be fulfilled by the lead contact, Lone Brøndsted.

### Materials availability

Bacterial isolates and bacteriophages are available by request from the lead contact under the conditions of a material transfer agreement (MTA).

### Data and code availability

Assembled bacteriophage genomes have been deposited at NCBI and are publicly available as of the date of publication. For accession numbers for phage genomes see Table 1. Bacterial genomes are available at NCBI under designated Bioprojects as noted in Table S1. This paper does not report original code.

## EXPERIMENTAL MODEL AND SUBJECT DETAILS

### Bacterial strains

ESBL/AmpC *E. coli* strains (n=198) (Table S1) originating from poultry, broiler meat or pig caecum were collected as part of Danish surveillance program and previously characterized (Vitt et al., 2023). *E. coli* strains were cultured in Lysogeny Broth (LB) and LB agar (LA) (Oxoid, Roskilde, Denmark). LA plates were supplemented with cefotaxime (Sigma Aldrich, Copenhagen, Denmark) at final concentration of 1 µg/ml. Overnight cultures were prepared in LB with 2-3 colonies shaking at 200 rpm at 37°C for 16-20 hours. Among 198 ESBL/AmpC *E. coli*, 19 strains were chosen for the isolation of the phages based on their characteristics as summarized in Figure 1.

## METHODS DETAILS

### Processing of animal and wastewater samples

Samples were collected from broiler faeces (n=6), pig waste (n=4); wastewater (n=2) was used to isolate phages. Faeces were diluted (1:10 w/v) in sterile SM buffer (0.1 M NaCl, 8 mM MgSO_4_, 50 mM Tris-HCl, pH 7.5), vortexed and centrifuged at 10.000 x g for 10 min at room temperature. Waste and wastewater samples were centrifuged at 6000 x g for 10 min. Following centrifugation, all supernatants were filtered through 0.45 µm filters and stored at 4°C.

### Phage isolation, purification, and propagation

Depending on the size of the Petri dish used, the bacterial lawns were prepared from 100 or 300 µl overnight cultures of isolation strains that were mixed with 4 or 10 ml of molten overlay agar (LBov; LB broth with 0,6% Agar bacteriological no.1 (Oxoid)) and spread on 9 or 12 cm LA (LB with 1,2% agar) plates, respectively. Bacterial lawns were allowed to settle for 15 minutes and then dried in a laminar hood for 45 min to be used immediately thereafter. For phage isolation, a total of 5 drops of 10 µl of sample were spotted on the lawns of isolation host strains and were incubated ON at 37°C aerobically. When no plaques were detected, the samples were subjected to selective enrichment with the isolation strains: 500 µl filtered sample, 500 µl ON isolation host culture and 1 ml LB mixed and incubated ON at 37°C with shaking at 180 rpm. The following day, the enrichment inoculums were centrifuged at 10.000 x g for 10 min and ten-fold serial dilutions in SM buffer were spotted on a lawn of the enrichment strain. Up to three plaques with different and consistent plaque morphologies were picked with a pipette tip and suspended in 200-400 µl of SM buffer, vortexed and ten-fold diluted. A 100 µl aliquot of selected dilution(s) were mixed with 100 µl of the isolation strain in 4 ml LBov and spread on LA plates. Each single plaque was purified for at least three rounds. Single plaques from the final purification steps were used for phage propagation on the isolation strain and phage stocks were prepared by plate lysis method adapted from ^58^ with modifications. Briefly, 100 µl of predetermined phage dilution, corresponding to a confluent lysis, was mixed with 100 µl of ON inoculum of the host strain, prepared as described above. After 10 min, 4 ml molten LBov was added, mixed gently, and poured over a pre-made LA plate. After settling of the overlay agar, the plates were incubated ON at 37°C. The next day, plates were examined, the layer with overlay agar was scraped off with a sterile inoculation loop, collected into a centrifuge tube and mixed with 5 ml SM-buffer. After thorough vortexing, the mixture was centrifuged at 8000 x g for 10 min at 4°C and the supernatant filtered once through 0.22 µm filters and stored at 4°C.

### Phage plaque assay

To determine phage titers, ten-fold serial dilutions (up to 10^-7^ – 10^-8^) of the phage stocks in SM buffer were prepared and 3 droplets of 10 µl aliquots were spotted on pre-made plates of bacterial lawns. Following an overnight incubation at 37°C, plaques were counted and plaque forming units per ml (PFU ml^-1^) were calculated for each strain.

### Host range analysis

Phage host range was determined in two steps: spot assay and, if lysis spots were observed, plaque assay was performed. For the spot assay, bacterial lawns were prepared in 12 cm round plates as described above. Brain Heart Infusion (BHI) (Oxoid) with 1.2 % agar for the basal plates and 0.6% agar for the overlays were used throughout the host range experiments. 10 µl of ten-fold diluted phage stocks (titers above 10^8^ PFU ml^-1^) were spotted on the air-dried bacterial lawns prepared with BHI 1.2 and 0.6 % agar and incubated 18-24 h at room temperature. To confirm phage infection for the spots with lysis, plaque assay was performed with one spot of 10 µl of each dilution. The plates were incubated 18-24 h at room temperature, plaques were counted and plaque forming units (PFU) ml^-1^ were calculated.

### Determination of receptors in *E. coli* strain ECOR4

Specific gene knockout strains were obtained with the Quick & Easy *E. coli* Gene Deletion Kit (Gene Bridges). Linear gene deletion fragments were generated by PCR (Amplicon Taq 2× Master Mix Red) using primers designed to match the FRT-PGK-gb2-neo-FRT cassette supplied with the mutagenesis kit. Each of the primer pairs were equipped with 50 bp 5’ extensions matching the terminal nucleotides of the respective genes of the ECOR4 chromosome. PCR products were purified and concentrated using the GeneJET PCR Purification Kit (Thermo Fischer Scientific) by elution in 10 µl nuclease-free water. First, *E. coli* ECOR4 were made competent by inoculation of 1 ml of an overnight culture in 100 ml fresh LB and incubation for 2.5 hours at 37°C with shaking at 180 rpm. The cells were put on ice for 10 min and collected by centrifugation at 4°C for 3 min at 4500 x g followed by two washes in 20 ml ice-cold 0.1 M CaCl_2_ before resuspension in 5 ml ice-cold 0.1 M CaCl_2_. 100 µl of competent cells were incubated on ice with 1 µl purified pRedET (amp) plasmid for 30 min before heat shock for 1 min at 42°C and addition of 900 µl LB followed by incubation for 1 hour at 30°C with shaking at 180 rpm. Cells were plated on LA containing 100 µg/ml ampicillin and incubated overnight at 30°C. Next, ECOR4 bearing the Red/ET expression plasmid were grown in a shaking flask in LB plus 100 µg/ml ampicillin at 30°C until an OD_600_ of 0.3 followed by addition of L-arabinose at 0.35% final concentration and continued growth at 37°C for 1 hour. Cells were cooled on ice and washed four times in ice-cold water (2 times 1 volume, 1 time ½ the volume, and 1 time ¼ the volume) and gently resuspended in 1/100 the volume of ice-cold water. 50 µl of cells were added 2 µl of concentrated deletion fragment and electroporated in pre-chilled 0.2 cm electroporation cuvettes using a MicroPulser Electroporator (Bio-Rad) at Ec2 settings. Fresh LB was added, and cells were incubated at 37°C for 2-3 hours before plating on LA containing 50 µg/ml kanamycin and overnight incubation at 37°C. Successful gene knockout was confirmed with gene-specific primers for respective target genes. After the construction of the ECOR4 mutants, the phage infectivity was evaluated using a normal plaque assay (see method above).

### Single bacteriophage and bacteriophage cocktail growth inhibition assay

To evaluate the phage growth inhibition effect of ESBL/AmpC *E. coli* strains, a phage cocktail was designed. ESBL/AmpC *E. coli* strains ESBL102 and ESBL145 was chosen as hosts because of their genetic diversity. Phages AV114, AV118 and AV125 were chosen as phage cocktail against ESBL102 whereas phages AV110, AV111, AV114 and AV125 were combined against ESBL145. Both single phages and the cocktail were evaluated for their growth inhibition ability against the strains. A single colony of each strain were inoculated in 5 mL LB media and incubated overnight at 37°C and 180 rpm. The following morning, the cultures were diluted to 1*10^8 CFU/mL and 100 µL of the culture was added to 96 wells plates (TPP). Afterwards, the phages (MOI of 10) were added to the samples and the plates were incubated in Gen5 plate reader (Agilent BioTek) for 24 hours at 37°C and OD600 values were determinate every 15 minutes. ESBL102 and ESBL145 without phages were used as negative controls. The growth inhibition assay was done in three triplicates and the standard deviations were calculated in GraphPad Prism9 (version 9.5.0)

### DNA extraction and sequencing

High titer (>10^8^ PFU ml^-1^) phage stocks were subjected to DNA extraction and purification by ethanol precipitation with modifications ^59^. Briefly, phage stocks were treated with RNase A (Thermo Fischer Scientific, Waltham, MA, USA) and DNAse I (Thermo Fischer Scientific) at final concentrations of 10 and 20 µg ml^-1^ and incubated at 37°C for 1-2 h. Phage DNA was released from capsids by treatment with proteinase K (50 µg/ml, Thermo Fischer Scientific) in the presence of SDS (0.5%) at 56°C overnight. After cooling the samples to room temperature, DNA was precipitated with 0.1 volume of 3 M sodium acetate (pH 5.5), glycogen (final concentration of 0.05 µg/µl, Thermo Fischer Scientific) and 2.5 volumes of ice-cold ethanol (96%) were added and incubated 2-6 days at –20°C. Precipitated DNA was centrifuged at 10000rpm for 30 min and washed two times with 70% ice-cold ethanol (10000rpm, 20 min). Pellets with DNA were air-dried at 37°C and dissolved in 10 mM Tris-HCl (pH 8) at 4°C overnight. Dissoved DNA was further purified using DNA Clean & Concentrator-25 (Zymo Research) following manufacturer’s instructions with elution in 50-150 µl of 10 mM Tris-HCl (pH 8). DNA concentrations were measured using Qubit (Thermo Fischer Scientific) and DNA libraries were prepared using Nextera XT v.3 (Illumina, San Diego, CA, USA) kit. Next generation sequencing was performed using MiSeq (Illumina) platform with paired-end (2 X 250-bp) mode.

## QUANTIFICATION AND STATISTICAL ANALYSIS

### Genome assembly and analyses

The sequences were *de novo* assembled using CLC Genomics Workbench 9.5.3 (Qiagen Digital Insights, Aarhus, Denmark). Open reading frames and tRNAs were detected and annotated automatically using Rapid Annotation Subsystem Technology RAST version 2.0 ^60^. tRNAs were additionally determined with ARAGORN software ^61^. The phages were assigned their taxonomy by overall genome BLAST similarities to their closest phage genome available at the NCBI.

### Bioinformatics analyses

The 19 isolating ESBL/AmpC E. coli genomes was extracted from Enterobase and a phylogenetic tree based on their cgMLST was visualized in iTOL v. 6 ^57^. CLC Workbench version 22 (Qiagen Digital Insights, Aarhus, Denmark) was used to alignment of all phage genomes with the default settings (minimum initial seed length: 15, allow mismatches in seeds: yes and minimum alignment block length: 100). Pairwise comparison of the analysis was conducted to create a heatmap displaying the average nucleotide identity (ANI) using default settings (table types: ANI, distance measure: euclidean distance and linkage criteria: complete linkage). Easyfig version 2.2.5 with 0.4 minimum identity for BLAST setting was used to align and visualize the phage genomes in the *Tequatrovirus*, *Mosigvirus* and *Phapecoctavirus* genera (Sullivan, Petty, and Beatson 2011). Furthermore, ClustalO ^62^ available in CLC was used for alignment of receptor binding protein sequences (Gp37 and Gp38) of the phages in the *Tevenvirnae* subfamily with default settings; Gap cost 10, gap extension cost 1, end gap cost: as any others and alignment mode: very accurate. VirulenceFinder version 2 ^36^ was used to estimate if any virulence genes were present in the phage genomes. The Gp37 of phage T4 (Accession number MT984581) and Gp38 of phage T2 (Accession number MH751506) and T6 (Accession number AP018814) was used for alignment of the Gp37 and Gp38 of Tevenvirinae phages isolated in the study.

