## Supplementary information for "A collection of diverse bacteriophages for biocontrol of ESBL- and AmpC-β-lactamase-producing *E. coli*"

**Supplementary Table S1:** Access to ESBL/AmpC *E. coli* sequences (Vitt et al. 2023)

**Supplementary Table S2:** The ability of each phage to infect diverse phylogroup and ST

**Supplementary Table S3:** Host range data of all phages on all ESBL/AmpC *E. coli* strains with pfu per ml (will be supplied as an Excel file)

**Supplementary Figure S1:** Protein sequence alignment of gp38 corresponding to the chaperone encoded by T4.

**Supplementary Figure S2.** Protein sequence alignment of the tail fibers corresponding to the adhesins encoded by gp38 homologues of the phages T2 and T6.

**Supplementary Figure S3.** Protein sequence alignment of gp37 of T4 and its homologues.

### Supplementary information

**Table S1: Access to ESBL/AmpC *E. coli* sequences (Vitt et al. 2023).**

| Strain name | Study Accession | Sample Accession | Secondary Sample Accession |
|---|---|---|---|
| ESBL1 | PRJEB22091 | SAMEA104205957 | ERS1864975 |
| ESBL2 | PRJEB22091 | SAMEA104205964 | ERS1864982 |
| ESBL3 | PRJEB22091 | SAMEA104205966 | ERS1864984 |
| ESBL4 | PRJEB22091 | SAMEA104205968 | ERS1864986 |
| ESBL5 | PRJEB22091 | SAMEA104205969 | ERS1864987 |
| ESBL6 | PRJEB22091 | SAMEA104205974 | ERS1864992 |
| ESBL7 | PRJEB22091 | SAMEA104205976 | ERS1864994 |
| ESBL8 | PRJEB22091 | SAMEA104205977 | ERS1864995 |
| ESBL9 | PRJEB22091 | SAMEA104205980 | ERS1864998 |
| ESBL10 | PRJEB22091 | SAMEA104205984 | ERS1865002 |
| ESBL11 | PRJEB22091 | SAMEA104205985 | ERS1865003 |
| ESBL12 | PRJEB22091 | SAMEA104205986 | ERS1865004 |
| ESBL13 | PRJEB14641 | SAMEA4058476 | ERS1229586 |
| ESBL14 | PRJEB14641 | SAMEA4058477 | ERS1229587 |
| ESBL15 | PRJEB14641 | SAMEA4058478 | ERS1229588 |
| ESBL16 | PRJEB14641 | SAMEA4058461 | ERS1229571 |
| ESBL17 | PRJEB14086 | SAMEA3993565 | ERS1164675 |
| ESBL18 | PRJEB14086 | SAMEA3993566 | ERS1164676 |
| ESBL19 | PRJEB14086 | SAMEA3993567 | ERS1164677 |
| ESBL20 | PRJEB14086 | SAMEA3993568 | ERS1164678 |
| ESBL21 | PRJEB14086 | SAMEA3993569 | ERS1164679 |
| ESBL22 | PRJEB14086 | SAMEA3993570 | ERS1164680 |
| ESBL23 | PRJEB14086 | SAMEA3993571 | ERS1164681 |
| ESBL24 | PRJEB14086 | SAMEA3993572 | ERS1164682 |
| ESBL25 | PRJEB14086 | SAMEA3993573 | ERS1164683 |
| ESBL26 | PRJEB14086 | SAMEA3993574 | ERS1164684 |

| | | | |
|---|---|---|---|
| ESBL27 | PRJEB14086 | SAMEA3993575 | ERS1164685 |
| ESBL28 | PRJEB14086 | SAMEA3993576 | ERS1164686 |
| ESBL29 | PRJEB14086 | SAMEA3993577 | ERS1164687 |
| ESBL30 | PRJEB14086 | SAMEA3993578 | ERS1164688 |
| ESBL31 | PRJEB14086 | SAMEA3993581 | ERS1164691 |
| ESBL32 | PRJEB14086 | SAMEA3993582 | ERS1164692 |
| ESBL33 | PRJEB14086 | SAMEA3993583 | ERS1164693 |
| ESBL34 | PRJEB14086 | SAMEA3993584 | ERS1164694 |
| ESBL35 | PRJEB14086 | SAMEA3993585 | ERS1164695 |
| ESBL36 | PRJEB14086 | SAMEA3993587 | ERS1164697 |
| ESBL37 | PRJEB14086 | SAMEA3993588 | ERS1164698 |
| ESBL38 | PRJEB14086 | SAMEA3993589 | ERS1164699 |
| ESBL39 | PRJEB14086 | SAMEA3993591 | ERS1164701 |
| ESBL40 | PRJEB14086 | SAMEA3993592 | ERS1164702 |
| ESBL41 | PRJEB14086 | SAMEA3993593 | ERS1164703 |
| ESBL42 | PRJEB14086 | SAMEA3993594 | ERS1164704 |
| ESBL43 | PRJEB14086 | SAMEA3993595 | ERS1164705 |
| ESBL44 | PRJEB14086 | SAMEA3993596 | ERS1164706 |
| ESBL45 | PRJEB14086 | SAMEA3993598 | ERS1164708 |
| ESBL46 | PRJEB14086 | SAMEA3993600 | ERS1164710 |
| ESBL47 | PRJEB14086 | SAMEA3993606 | ERS1164716 |
| ESBL48 | PRJEB14086 | SAMEA3993607 | ERS1164717 |
| ESBL49 | PRJEB14086 | SAMEA3993608 | ERS1164718 |
| ESBL50 | PRJEB14086 | SAMEA3993609 | ERS1164719 |
| ESBL51 | PRJEB14086 | SAMEA3993610 | ERS1164720 |
| ESBL52 | PRJEB14086 | SAMEA3993611 | ERS1164721 |
| ESBL53 | PRJEB14086 | SAMEA3993612 | ERS1164722 |
| ESBL54 | PRJEB14086 | SAMEA3993615 | ERS1164725 |
| ESBL55 | PRJEB14086 | SAMEA3993616 | ERS1164726 |

| | | | |
|---|---|---|---|
| ESBL56 | PRJEB14086 | SAMEA3993617 | ERS1164727 |
| ESBL57 | PRJEB14641 | SAMEA4058361 | ERS1229471 |
| ESBL58 | PRJEB14641 | SAMEA4058362 | ERS1229472 |
| ESBL59 | PRJEB14641 | SAMEA4058363 | ERS1229473 |
| ESBL60 | PRJEB14641 | SAMEA4058365 | ERS1229475 |
| ESBL61 | PRJEB14641 | SAMEA4058366 | ERS1229476 |
| ESBL62 | PRJEB14641 | SAMEA4058367 | ERS1229477 |
| ESBL63 | PRJEB14641 | SAMEA4058368 | ERS1229478 |
| ESBL64 | PRJEB14641 | SAMEA4058371 | ERS1229481 |
| ESBL65 | PRJEB14641 | SAMEA4058372 | ERS1229482 |
| ESBL66 | PRJEB14641 | SAMEA4058373 | ERS1229483 |
| ESBL67 | PRJEB14641 | SAMEA4058374 | ERS1229484 |
| ESBL68 | PRJEB14641 | SAMEA4058377 | ERS1229487 |
| ESBL69 | PRJEB14641 | SAMEA4058379 | ERS1229489 |
| ESBL70 | PRJEB14641 | SAMEA4058380 | ERS1229490 |
| ESBL71 | PRJEB14641 | SAMEA4058381 | ERS1229491 |
| ESBL72 | PRJEB14641 | SAMEA4058382 | ERS1229492 |
| ESBL73 | PRJEB14641 | SAMEA4058384 | ERS1229494 |
| ESBL74 | PRJEB14641 | SAMEA4058385 | ERS1229495 |
| ESBL75 | PRJEB14641 | SAMEA4058386 | ERS1229496 |
| ESBL76 | PRJEB14641 | SAMEA4058387 | ERS1229497 |
| ESBL77 | PRJEB14641 | SAMEA4058388 | ERS1229498 |
| ESBL78 | PRJEB14641 | SAMEA4058389 | ERS1229499 |
| ESBL79 | PRJEB14641 | SAMEA4058390 | ERS1229500 |
| ESBL80 | PRJEB14641 | SAMEA4058391 | ERS1229501 |
| ESBL81 | PRJEB14641 | SAMEA4058392 | ERS1229502 |
| ESBL82 | PRJEB14641 | SAMEA4058393 | ERS1229503 |
| ESBL83 | PRJEB14641 | SAMEA4058395 | ERS1229505 |
| ESBL84 | PRJEB14641 | SAMEA4058397 | ERS1229507 |

| | | | |
|---|---|---|---|
| ESBL85 | PRJEB14641 | SAMEA4058400 | ERS1229510 |
| ESBL86 | PRJEB14641 | SAMEA4058401 | ERS1229511 |
| ESBL87 | PRJEB14641 | SAMEA4058402 | ERS1229512 |
| ESBL88 | PRJEB14641 | SAMEA4058403 | ERS1229513 |
| ESBL89 | PRJEB14641 | SAMEA4058404 | ERS1229514 |
| ESBL90 | PRJEB14641 | SAMEA4058405 | ERS1229515 |
| ESBL91 | PRJEB14641 | SAMEA4058406 | ERS1229516 |
| ESBL92 | PRJEB14641 | SAMEA4058407 | ERS1229517 |
| ESBL93 | PRJEB14641 | SAMEA4058408 | ERS1229518 |
| ESBL94 | PRJEB14641 | SAMEA4058409 | ERS1229519 |
| ESBL95 | PRJEB14641 | SAMEA4058410 | ERS1229520 |
| ESBL96 | PRJEB14641 | SAMEA4058411 | ERS1229521 |
| ESBL97 | PRJEB14641 | SAMEA4058412 | ERS1229522 |
| ESBL98 | PRJEB14641 | SAMEA4058413 | ERS1229523 |
| ESBL99 | PRJEB14641 | SAMEA4058414 | ERS1229524 |
| ESBL100 | PRJEB14641 | SAMEA4058415 | ERS1229525 |
| ESBL101 | PRJEB14641 | SAMEA4058416 | ERS1229526 |
| ESBL102 | PRJEB14641 | SAMEA4058417 | ERS1229527 |
| ESBL103 | PRJEB14641 | SAMEA4058418 | ERS1229528 |
| ESBL104 | PRJEB14641 | SAMEA4058419 | ERS1229529 |
| ESBL105 | PRJEB14641 | SAMEA4058420 | ERS1229530 |
| ESBL106 | PRJEB14641 | SAMEA4058421 | ERS1229531 |
| ESBL107 | PRJEB14641 | SAMEA4058422 | ERS1229532 |
| ESBL108 | PRJEB14641 | SAMEA4058423 | ERS1229533 |
| ESBL109 | PRJEB14641 | SAMEA4058425 | ERS1229535 |
| ESBL110 | PRJEB14641 | SAMEA4058426 | ERS1229536 |
| ESBL111 | PRJEB14641 | SAMEA4058427 | ERS1229537 |
| ESBL112 | PRJEB14641 | SAMEA4058428 | ERS1229538 |
| ESBL113 | PRJEB14641 | SAMEA4058429 | ERS1229539 |

| | | | |
|---|---|---|---|
| ESBL114 | PRJEB14641 | SAMEA4058430 | ERS1229540 |
| ESBL115 | PRJEB14641 | SAMEA4058431 | ERS1229541 |
| ESBL116 | PRJEB14641 | SAMEA4058432 | ERS1229542 |
| ESBL117 | PRJEB14641 | SAMEA4058433 | ERS1229543 |
| ESBL118 | PRJEB14641 | SAMEA4058434 | ERS1229544 |
| ESBL119 | PRJEB14641 | SAMEA4058435 | ERS1229545 |
| ESBL120 | PRJEB14641 | SAMEA4058436 | ERS1229546 |
| ESBL121 | PRJEB14641 | SAMEA4058437 | ERS1229547 |
| ESBL122 | PRJEB14641 | SAMEA4058438 | ERS1229548 |
| ESBL123 | PRJEB14641 | SAMEA4058439 | ERS1229549 |
| ESBL124 | PRJEB14641 | SAMEA4058440 | ERS1229550 |
| ESBL125 | PRJEB14641 | SAMEA4058441 | ERS1229551 |
| ESBL126 | PRJEB14641 | SAMEA4058442 | ERS1229552 |
| ESBL127 | PRJEB14641 | SAMEA4058468 | ERS1229578 |
| ESBL128 | PRJEB14641 | SAMEA4058469 | ERS1229579 |
| ESBL129 | PRJEB14641 | SAMEA4058470 | ERS1229580 |
| ESBL130 | PRJEB14641 | SAMEA4058471 | ERS1229581 |
| ESBL131 | PRJEB14641 | SAMEA4058472 | ERS1229582 |
| ESBL132 | PRJEB14641 | SAMEA4058473 | ERS1229583 |
| ESBL133 | PRJEB22091 | SAMEA104205989 | ERS1865007 |
| ESBL134 | PRJEB22091 | SAMEA104205991 | ERS1865009 |
| ESBL135 | PRJEB22091 | SAMEA104205992 | ERS1865010 |
| ESBL136 | PRJEB22091 | SAMEA104205993 | ERS1865011 |
| ESBL137 | PRJEB22091 | SAMEA104205996 | ERS1865014 |
| ESBL138 | PRJEB22091 | SAMEA104205998 | ERS1865016 |
| ESBL139 | PRJEB22091 | SAMEA104205999 | ERS1865017 |
| ESBL140 | PRJEB22091 | SAMEA104206001 | ERS1865019 |
| ESBL141 | PRJEB22091 | SAMEA104206002 | ERS1865020 |
| ESBL142 | PRJEB22091 | SAMEA104206003 | ERS1865021 |

| | | | |
|---|---|---|---|
| ESBL143 | PRJEB22091 | SAMEA104206004 | ERS1865022 |
| ESBL144 | PRJEB22091 | SAMEA104206005 | ERS1865023 |
| ESBL145 | PRJEB22091 | SAMEA104206006 | ERS1865024 |
| ESBL146 | PRJEB22091 | SAMEA104206007 | ERS1865025 |
| ESBL147 | PRJEB22091 | SAMEA104206008 | ERS1865026 |
| ESBL148 | PRJEB22091 | SAMEA104206009 | ERS1865027 |
| ESBL149 | PRJEB22091 | SAMEA104206010 | ERS1865028 |
| ESBL150 | PRJEB22091 | SAMEA104206011 | ERS1865029 |
| ESBL151 | PRJEB22091 | SAMEA104206012 | ERS1865030 |
| ESBL152 | PRJEB22091 | SAMEA104206013 | ERS1865031 |
| ESBL153 | PRJEB22091 | SAMEA104206014 | ERS1865032 |
| ESBL154 | PRJEB22091 | SAMEA104206015 | ERS1865033 |
| ESBL155 | PRJEB22091 | SAMEA104206017 | ERS1865035 |
| ESBL156 | PRJEB22091 | SAMEA104206018 | ERS1865036 |
| ESBL157 | PRJEB22091 | SAMEA104206019 | ERS1865037 |
| ESBL158 | PRJEB22091 | SAMEA104206020 | ERS1865038 |
| ESBL159 | PRJEB22091 | SAMEA104206021 | ERS1865039 |
| ESBL160 | PRJEB22091 | SAMEA104205896 | ERS1864914 |
| ESBL161 | PRJEB22091 | SAMEA104205900 | ERS1864918 |
| ESBL162 | PRJEB22091 | SAMEA104205901 | ERS1864919 |
| ESBL163 | PRJEB22091 | SAMEA104205902 | ERS1864920 |
| ESBL164 | PRJEB22091 | SAMEA104205903 | ERS1864921 |
| ESBL165 | PRJEB22091 | SAMEA104205904 | ERS1864922 |
| ESBL166 | PRJEB22091 | SAMEA104205906 | ERS1864924 |
| ESBL167 | PRJEB22091 | SAMEA104205907 | ERS1864925 |
| ESBL168 | PRJEB22091 | SAMEA104205910 | ERS1864928 |
| ESBL169 | PRJEB22091 | SAMEA104205912 | ERS1864930 |
| ESBL170 | PRJEB22091 | SAMEA104205913 | ERS1864931 |
| ESBL171 | PRJEB22091 | SAMEA104205915 | ERS1864933 |

| | | | |
|---|---|---|---|
| ESBL172 | PRJEB22091 | SAMEA104205916 | ERS1864934 |
| ESBL173 | PRJEB22091 | SAMEA104205917 | ERS1864935 |
| ESBL174 | PRJEB22091 | SAMEA104205918 | ERS1864936 |
| ESBL175 | PRJEB22091 | SAMEA104205919 | ERS1864937 |
| ESBL176 | PRJEB22091 | SAMEA104205920 | ERS1864938 |
| ESBL177 | PRJEB22091 | SAMEA104205921 | ERS1864939 |
| ESBL178 | PRJEB22091 | SAMEA104205922 | ERS1864940 |
| ESBL179 | PRJEB22091 | SAMEA104205923 | ERS1864941 |
| ESBL180 | PRJEB22091 | SAMEA104205924 | ERS1864942 |
| ESBL181 | PRJEB22091 | SAMEA104205925 | ERS1864943 |
| ESBL182 | PRJEB22091 | SAMEA104205927 | ERS1864945 |
| ESBL183 | PRJEB22091 | SAMEA104205930 | ERS1864948 |
| ESBL184 | PRJEB22091 | SAMEA104205932 | ERS1864950 |
| ESBL185 | PRJEB22091 | SAMEA104205934 | ERS1864952 |
| ESBL186 | PRJEB22091 | SAMEA104205936 | ERS1864954 |
| ESBL187 | PRJEB22091 | SAMEA104205937 | ERS1864955 |
| ESBL188 | PRJEB22091 | SAMEA104205939 | ERS1864957 |
| ESBL189 | PRJEB22091 | SAMEA104205941 | ERS1864959 |
| ESBL190 | PRJEB22091 | SAMEA104205942 | ERS1864960 |
| ESBL191 | PRJEB22091 | SAMEA104205943 | ERS1864961 |
| ESBL192 | PRJEB22091 | SAMEA104205945 | ERS1864963 |
| ESBL193 | PRJEB22091 | SAMEA104205949 | ERS1864967 |
| ESBL194 | PRJEB22091 | SAMEA104205951 | ERS1864969 |
| ESBL195 | PRJEB22091 | SAMEA104205953 | ERS1864971 |
| ESBL196 | PRJEB22091 | SAMEA104205954 | ERS1864972 |
| ESBL197 | PRJEB22091 | SAMEA104205955 | ERS1864973 |
| ESBL198 | PRJEB22091 | SAMEA104205956 | ERS1864974 |

**Supplementary Table S2.** The ability of each phage to infect diverse phylogroup and STs

| Phage | Number of infected hosts | Phylogroups | STs based on MLST analysis |
|---|---|---|---|
| AV101 | 9 | A, B1, C | ST-1800, ST-2040, ST-6206, ST-641, ST-88 |
| AV102 | 4 | B1, C | ST-162, ST-1431, ST-23 |
| AV103 | 4 | B1, C | ST-23, ST-641, ST-6254, ST-453 |
| AV104 | 2 | D | ST-69, ST-4243 |
| AV105 | 2 | D | ST-69, ST-4243 |
| AV106 | 1 | C | ST88 |
| AV107 | 14 | A, B1, C, D | ST-23, ST-4980, ST-4243, ST-101 |
| AV108 | 5 | A, B1 | ST-295, ST-10, ST-154, ST-6206 |
| AV109 | 9 | B1, C | ST-88, ST-7614, ST-4663, ST-295 |
| AV110 | 48 | A, B1, B2, C, G | ST-10, ST-101, ST-117, ST-1431, ST-154, ST-155, ST-156, ST-23, ST-295, ST-345, ST-3564, ST-429, ST-4580, ST-4663, ST-4980, ST-58, ST-665, ST-718, ST-75, ST-88 |
| AV111 | 46 | A, B1, C, D, E, F | ST-10, ST-1147, ST-1640, ST-23, ST-2607, ST-354, ST-4243, ST-453, ST-4663, ST-4980, ST-57, ST-58, ST-641 |
| AV112 | 37 | A, B1, C, D | ST-10, ST-101, ST-1011, ST-1431, ST-154, ST-155, ST-156, ST-1800, ST-2040, ST-295, ST-3564, ST-4580, ST-48, ST-4980, ST-58, ST-718, ST-75, ST-877, ST-88 |
| AV113 | 25 | A, B1, C, E | ST-295, ST-10, ST-101, ST-1431, ST-154, ST-155, ST-156, ST-1800, ST-295, ST-3564, ST-48, ST-57, ST-58, ST-718, ST-877, ST-88 |
| AV114 | 29 | A, B1, C, G | ST-117, ST-155, ST-165, ST-295, ST-3564, ST-453, ST-4663, ST-4980, ST-718, ST-7614, ST-88 |
| AV115 | 33 | A, B1, C, G | ST-101, ST-117, ST-1431, ST-165, ST-295, ST-3564, ST-453, ST-4663, ST-58, ST-88 |
| AV116 | 17 | A, B1, C | ST-101, ST-1431, ST-154, ST-155, ST-156, ST-1800, ST-2040, ST-295, ST-4580, ST-6206, ST-88 |
| AV117 | 16 | A, B1, C | ST-101, ST-1431, ST-154, ST-155, ST-156, ST-1800, ST-2040, ST-295, ST-4580, ST-6206, ST-88 |
| AV118 | 9 | A, B1, C | ST-155, ST-1850, ST-4580, ST-88 |
| AV119 | 55 | A, B1, B2, C, D, E, F | ST-10, ST-1147, ST-115, ST-131, ST-1640, ST-23, ST-355, ST-367, ST-4243, ST-453, ST-4663, ST-4980, ST-57, ST-6858, ST-69, ST-877, ST-88 |

| | | | |
|---|---|---|---|
| AV120 | 50 | A, B1, B2, C, D, E | ST-10, ST-131, ST-1640, ST-23, ST-2607, ST-367, ST-4243, ST-453, ST-4663, ST-4980, ST-57, ST-58, ST-602, ST-877, ST-88 |
| AV121 | 26 | Clade I, A, B1, B2, C, D | ST-115, ST-131, ST-2309, ST-355, ST-4243, ST-453, ST-4980, ST-770, ST-88 |
| AV122 | 16 | Clade I, B1, C, E | ST-101, ST-1431, ST-154, ST-155, ST-1640, ST-4580, ST-770, ST-877, ST-88 |
| AV123 | 51 | A, B1, B2, C, D, E | ST-101, ST-1056, ST-1147, ST-1286, ST-131, ST-1463, ST-154, ST-155, ST-162, ST-1800, ST-1850, ST-2040, ST-23, ST-2607, ST-38, ST-4243, ST-429, ST-4580, ST-4663, ST-4980, ST-57, ST-602, ST-6206, ST-718, ST-88 |
| AV124 | 2 | C | ST-23, ST-88 |
| AV125 | 84 | A, B1, B2, C, D, E | ST-10, ST-101, ST-1011, ST-1056, ST-1147, ST-115, ST-1431, ST-1463, ST-154, ST-155, ST-156, ST-162, ST-1640, ST-1800, ST-1850, ST-2040, ST-23, ST-2309, ST-2607, ST-295, ST-345, ST-3564, ST-367, ST-38, ST-4243, ST-429, ST-4580, ST-4663, ST-4980, ST-57, ST-602, ST-6206, ST-6254, ST-718, ST-88 |
| AV126 | 65 | Clade I, A, B1, B2, C, D, E | ST-770, ST-10, ST-101, ST-1011, ST-115, ST-1463, ST-154, ST-155, ST-1640, ST-1800, ST-1850, ST-2040, ST-23, ST-2607, ST-295, ST-345, ST-351, ST-38, ST-4243, ST-429, ST-4580, ST-4980, ST-57, ST-602, ST-6206, ST-718, ST-877, ST-88 |
| AV127 | 13 | A, B1, C | ST-1463, ST-155, ST-23, ST-4980, ST-4980, ST-57, ST-602, ST-88 |
| AV128 | 8 | A, B1, D, E | ST-101, ST-1640, ST-2197, ST-38, ST-57 |
| AV129 | 12 | A, B1, C, D, E | ST-1640, ST-23, ST-2952, ST-4243, ST-4980 |

**Figure S1:** Protein sequence alignment of gp38 corresponding to the chaperone encoded by T4.

Differing residues are highlighted in red colour.

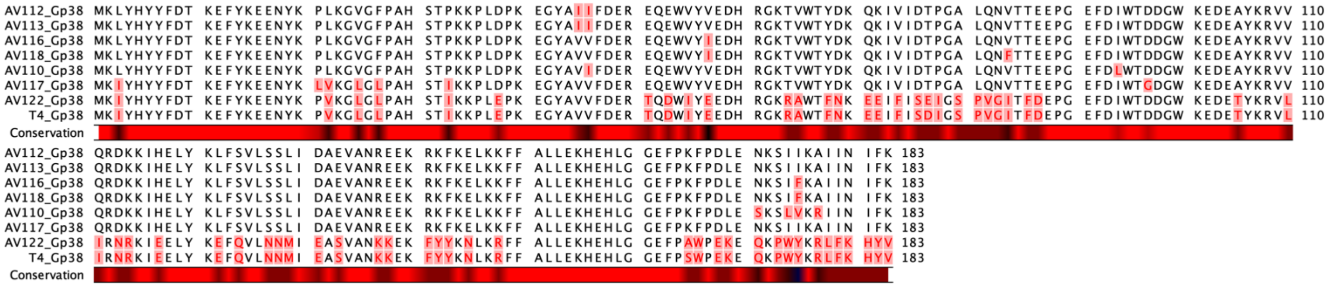

**Supplementary Figure S2.** Protein sequence alignment of the tail fibers corresponding to the adhesins encoded by gp38 homologues of the phages T2 and T6. Differing residues are highlighted in red colour.

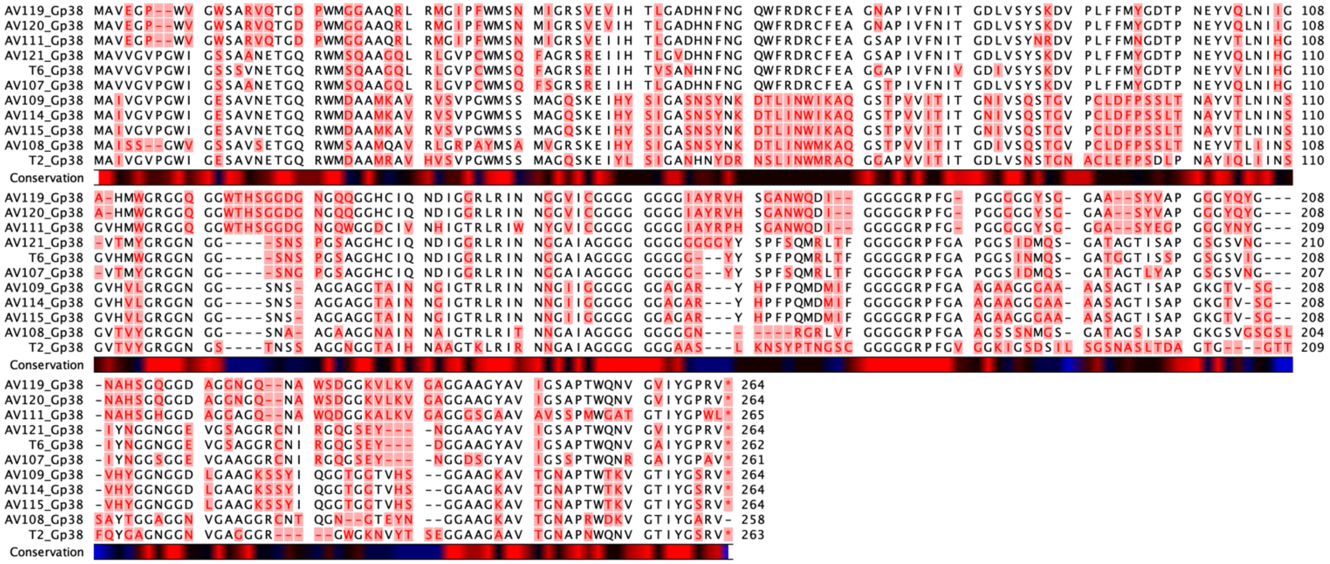

**Figure S3.** Protein sequence alignment of gp37 of T4 and its homologues. Alignment of full sequences of gp37 of phage T4 and AV110, AV112, AV113, AV116, AV117, AV118, AV122. Differing residues are highlighted in red.

```
AV110_Gp37 MATLKQIQFK RSKVAGVNPT AAQLAEGELA INLKDRTLFT KDDLGNIISL GFAKGGDING NVTQTGDYLQ NG------TY NLNGNQFVYA GKYIEFLPKT 94
AV118_Gp37 MATLKQIQFK RSKVAGVNPT AAQLAEGELA INLKDRTLFT KDDLGNIISL GFAKGGDING NVTQTGDYLQ NG------TY NLNGNQFVYA GKYIEFLPKT 94
AV112_Gp37 MATLKQIQFK RSKVAGVNPT AAQLAEGELA INLKDRTLFT KDDLGNIISL GFAKGGDING NVTQNGDYLQ NG------TY NLNGNQFVYA GKYIEFLPKT 94
AV113_Gp37 MATLKQIQFK RSKVAGVNPT AAQLAEGELA INLKDRTLFT KDDLGNIISL GFAKGGDING NVTQNGDYLQ NG------TY NLNGNQFVYA GKYIEFLPKT 94
AV117_Gp37 MATLKQIQFK RSKVAGVNPT AAQLAEGELA INLKDRTLFT KDDLGNIISL GFAKGGDING NVTQNGDYLQ NG------AY NLNGNQFVYA GKYIEFLPKT 94
   T4_Gp37 MATLKQIQFK RSKIAGTRPA ASVLAEGELA INLKDRTIFT KDDSGNIIDL GFAKGGQVDG NVTINGLLRL NGDYVQTGGM TVNGPIGSTD G-VTAKIFRS 99
AV116_Gp37 MATLKQIQFK RSKTAGARPA ASVLAEGELA INLKDRTLFT KDDSGNIIDL SISAGGNISG NVTIDGTLRV NGPINNFGNF STSGQITAGS S-ISAQVFRA 99
AV122_Gp37 MATLKQIQFK RSKTAGQRPA ASVLAEGELA INLKDRTLFT KDDSGNIIDL SITAGGNISG NITQTGDYTQ TG------KF NLIGPQIVAS GGYIEFNYRT 94
Conservation

AV110_Gp37 TGNGAWAN-- -QHSNKAPIF TDLSSTTSVS EYHPLIKQRY KDGTFSAGTL VNEGSFKFHY IDETG--TSK YWTFNRNGNF QVDSGSLLVS NGYLVTQSNI 189
AV118_Gp37 TGNGAWAN-- -QHSNKAPIF TDLSSTTSVS EYHPLIKQRY KDGTFSAGTL VNEGSFKFHY IDETG--TSK YWTFNRNGNF QVDSGSLLVS NGYLVTQSNI 189
AV112_Gp37 AGNGAWAN-- -QHTNKAPIF TDLSSTTSTS EYHPLIKQRY KNGTFSAGTL VSEGSFKIHY IDETG--DSK YWTFNRNGSF IVDNGNIEVR AGNISASGNI 189
AV113_Gp37 AGNGAWAN-- -QHTNKAPIF TDLSSTTSTS EYHPLIKQRY KNGTFSAGTL VSEGSFKIHY IDETG--DSK YWTFNRNGSF IVDNGNIEVR AGNISASGNI 189
AV117_Gp37 AGNGAWAN-- -QHTNKAPIF TDLSSTTSTS EYHPLIKQRY KDGTFSVGTL VSEGSFRVHY IDSTAPDNSK HWAFNRNGNF IVDSGSIEVR AGNISASGNI 191
   T4_Gp37 TQGSFYARAT -NDTSNAHLW FENADGTER- ---GVI---- ----YARPQT TTDGEIRLRV RQGTGSTANS EFYFRS---- ---------IN GGEF-QANRI 173
AV116_Gp37 LQGSFYSRAS TGETANAHLW FENADGTER- ---GVI---- ----YARPQT TTDGEIRLRV RQGTGSTVNS EFYFRS---- ---------IN GGEF-QANRI 174
AV122_Gp37 TGSGAWSG-- -QHTAKAPIF VDLSSATSTS EYNPIIKQRF KDGTFSLGTL VSEGSLKIHY INESGDS--K YWTFRRDGGF TVDGGGLAVS GGSITTTGNI 189
Conservation

AV110_Gp37 ETKAGSLIGP KVITKNINVS TKSWDTNIDT AYDLVSLPTW QYTPTE--TD LDTNGINYLR KFRGHVSSAI FHEVADARSG KPQNISYWTG RNAEVYQWSY 287
AV118_Gp37 ETKAGSLIGP KVITKNINVS TKSWDTNIDT AYDLVSLPTW HYTPTE--TD LDTNGINYLR KFRGHVSSAI FHEVADARSG KPQNISYWTG RNAEVYQWSY 287
AV112_Gp37 NSASGVVSAP RVNTKNINLS SKTFIPNNDA GYDIVSLPTW IYNPPADGSD RDTNGLNYMR KMRASNTSSI FHEIVDCRKD KPQEISWWTG VGPSSFQWSF 289
AV113_Gp37 NSASGVVSAP RVNTKNINLS SKTFIPNNDA GYDIVSLPTW IYNPPADGSD RDTNGLNYMR KMRASNTSSI FHEIVDCRKD KPQEISWWTG VGPSSFQWSF 289
AV117_Gp37 NSANGIVSAP QVNTKTIVLD TKAFG----- QYDSQSLVQY VYPGT----- GEENGINYLR KVRAKSGGTI YHEIASAQTG KNDEISWWTG NSLTTKLMGL 281
   T4_Gp37 L-ASDSLVTK RIAVDTVIHD AKAFG----- QYDSHSLVNY VYPGT----- GETNGVNYLR KVRAKSGGTI YHEIVTAQTG LADEVSWWSG DTPVFKLYGI 262
AV116_Gp37 L-ASDSLVTK RIAADTVIHG GKAFG----- VYDTDSLVNY VYPGT----- GETNGVNYLR KVRAKSGGTM YHEIVTAQTD QADEVSWWTG NTPSFKLYGI 263
AV122_Gp37 A-ALGNITSP QINTKNIILD TKAFG----- QYDSQSLVNY VYPGT----- GEENGINYLR KVRSKSGGTI YHEICTAQTG LADEVSWWTG NTPLFKLYGI 278
Conservation

AV110_Gp37 TTDGRMSVAN SIGIGLADVG DQG-RYPLVS --F-GKGIAI GDNDTGIKWV RDGVIDLMAN GISSATILNN GINSTKKLMV GYSRDSGSTW DFPNNNQAMF 383
AV118_Gp37 TTDGRMSVAN SIGIGLADVG VQG-RYPLVS --F-GKGIAI GDNDTGIKWV RDGVIDLIAN GVSTGTINST GVNSTKKLTV GYSRDSGSTW DYPNTNQAMF 383
AV112_Gp37 DTTGRIAAGK SISVGLPDNG VNG-RYPLES -AI-SIGIAI GDNDTGIKWV RDGVIDFMAN GISSATILNN GINSTKKLMV GYSRDSGTTW DFPNNNQAMF 386
AV113_Gp37 DTTGRIAAGK SISVGLPDNG VNG-RYPLES -AI-SIGIAI GDNDTGIKWV RDGVIDFMAN GISSATILNN GINSTKKLMV GYSRDSGTTW DFPNNNQAMF 386
AV117_Gp37 RNDGAMVLRR SLAIGTITTD ENTNNYGSPT PMG-ERYIAL GDAATGLKYI KQGVYDLVGN WNSVASITPD SFRSTRKALF GRSEDQGGTW TMPGTNAAFL 380
   T4_Gp37 RDDGRMIIRN SLALGTFTTN FPSSDYGNVG VMG-DKYLVL GDTVTGLSYK KTGVFDLVGG GYSVASITPD SFRSTRKGIF GRSEDQGATW IMPGTNAALL 361
AV116_Gp37 RNDGRMIVRN SIAIGTVTAG YPSSDYGNGA ALGFGVYLAL GDNGTGLSYK KTGVYDLVGS GNSVASITPD SFRSTRKAIF GRSEDQGTTW IMPGTNAAFL 363
AV122_Gp37 RNDGRMIIRN SLAIGTVTEG YPSSDYGNTG VMG-DRYLAI GDNASGLKWV RTGVYDLMAN GISSATILNN GINSTKKLMV GYSRDSGSTW DFPNNNQAMF 377
Conservation

AV110_Gp37 TARTDVDGNN NGDGQTHIGY SSNAKMYHYF RGTGRMSVSM GDGLLVEPGI LDIKTGSNSL NLRADGTIAS VQTLKLNNGL YFNGDGTNAS LVFKAASGID 483
AV118_Gp37 SARTEVDGNN NGDGQTHIGY SSNAKMYHYF RGAGRMAISM GEGLIVEPGI LDIKTGSNSL NLRADGTIAS VQTLKLNNGL YFNGDGTNAS LIFKAASGID 483
AV112_Gp37 TARTDIDGNN NGDGQTHIGY SSNSKMYHYF RGTGRMAIGM AEGLIVEPGI LDIKTSSNTL SLRADGTVAS VQTLKLNNGL YFNGDGTNAS LVFKAASGID 486
AV113_Gp37 TARTDIDGNN NGDGQTHIGY SSNSKMYHYF RGTGRMAIGM AEGLIVEPGI LDIKTSSNTL SLRADGTVAS VQTLKLNNGL YFNGDGTNAS LVFKAASGID 486
AV117_Gp37 SVQTQADNNN AGDGRTHIGY NSGGKMSHYF RGTGQMNINT QQGMEINPGI LKLVTGANNV MFYADGGITS TKQLSIYNGV YLAANNSTAG LTFAPTTTID 480
   T4_Gp37 SVQTQADNNN AGDGQTHIGY NAGGKMNHYF RGTGQMNINT QQGMEINPGI LKLVTGSNNV QFYADGTISS IQPIKLDNEI FLTKSNNTAG LKFGAPSQVD 461
AV116_Gp37 SVQTQADENT AGDGQTHIGY NSGGKMSHYF RGKGQTNINT QEGMEVNPGI LKLVTGSNNV QFYADGTVSS IQRIKFDNGL VLTGA-RPDG IQLDAPTTTD 462
AV122_Gp37 TARTDVDGNN NGDGQTHIGY NSNSKMYHYF RGAGRMSVSM GDGLLVEPGI LDIKTGSNRV AIRADGTFDS TQRISMRTGL FLNGDDNTNA LSFKAPTGVD 477
Conservation

AV110_Gp37 GTKQIMWEGG TRAGQNKSYV TVKAWGNSFN ATGDKARETV FEVSDGQGYY FYSQRVAPTG SETTGPVVTQ FNGQVNARGS LITDGSLRVN GLSTLVGNVT 583
AV118_Gp37 GTKQIMWEGG TRAGQNKSYV TVKAWGNSFN ASGDRNRETV FEVSDGQGYY FYSQRVAPSG SETTGPVVTQ FNGQVNARGS LITDGSLRVN GLSTLVGNVT 583
AV112_Gp37 GTKQIMWEGG TRAGQNKSYV TVKAWGNSFN ASGDRTRETV FEVADGQGYY FYSQRVAPTG SETVGPVVTQ INGQLNARGS IITDGSLRVN GLSTLSGQVT 586
AV113_Gp37 GTKQIMWEGG TRAGQNKSYV TVKAWGNSFN ASGDRTRETV FEVADGQGYY FYSQRVAPTG SETVGPVVTQ INGQLNARGS IITDGSLRVN GLSTLSGQVT 586
AV117_Gp37 GTKQILWEGG KRAGQNKSYV TVKAWGNSFN ASGDRNRETV FEVADGQGYY FYSQRVAPSG SETTGPVVTQ FAGAVNARGS IITDGSFKVN GLSTLVGNVT 580
   T4_Gp37 GTRTIQWNGG TREGQNKNYV IIKAWGNSFN ATGDRSRETV FQVSDSQGYY FYAHRKAPTG DETIGRIEAQ FAGDVYAKGI I-ANGNFRVV GSSALAGNVT 560
AV116_Gp37 GTKTILWAGG TREGQNKNYV SIKAWGNAFN ASGDRSRETV FEVGDGQGYY FYAQRVAPAT GSTIGSIQLR IGGALLTSGG I-TS------ -----SGSIV 550
AV122_Gp37 GTKTINWDAG TRNGQNKNTV SIKAWGNAFN SVP-GIRETV FEVYDSQGYY FYSQRQRPEG SETVGPVVTQ FNGPVNANGS ISTAGNLKVD GLSTLNGNVT 576
Conservation

AV110_Gp37 MNNGLFVQGG ASITGQVKIG GTADAFRIYD AEYGAIFRRS EDKFYIIPTP QNAGESGGIS -NLRPFYITL SNGKVSMDHD VDIGGGRFKV EAAGT----- 677
AV118_Gp37 MNNGLFVQGG SSITGQVKIG GTADALRIWN AEYGAIFRRS EASLYIIPTP KDAGESGGIS -NLRPLIIEL NTGTVKMSHD VHLGDSGSGT GLLQV--SNS 680
AV112_Gp37 MDNGLVLSGG GSITGQVKIG NTADAFRIYN AEYGAIFRRS EASLYIIPTP KDAGESGGIS -NLRPLTIEL NTGTVKMSHD VHLGDSGSGT GLLQV--SNS 683
AV113_Gp37 MDNGLVLSGG GSITGQVKIG NTADAFRIYN AEYGAIFRRS EASLYIIPTP KDAGESGGIS -NLRPLTIEL NTGTVKMSHD VHLGDSGSGT GLLQV--SNS 683
AV117_Gp37 MSNGLNLTGG SSITGQVKIG GTADALRIWN AEYGAIFRRS EDKLYIIPTP QNAGETGGIS -NLRPFYIAL SNGKVSMDHD VDIGGGRFKV ETSGT----- 674
   T4_Gp37 MSNGLFVQGG SSITGQVKIG GTANALRIWN AEYGAIFRRS ESNFYIIPTN QNEGESGDIH SSLRPVRIGL NDGMVGL--- --------GR DSFIVDQNNA 649
AV116_Gp37 TESSLVANNG MSVNGQAKFG GTANALRIWN AEYGAIFRRS ETALHIIPTL KDQGENGEIG -NLRPLSISL HNGMVQMRHS VTLGDDGAGG NMITVDNDNK 649
AV122_Gp37 MKNGLFVQGG SSITGQVKIG GTTNALRIWN AEYGAIFRRS ESNFYIIPTN KNEGESGDIH SSLRPVRIGL NDGMVGL--- --------GR DSFIVDQNNA 665
Conservation

AV110_Gp37 ----TAGQRI TVNMASDA-- ---IRIN--- --APTQESSN FIQGRKADVP KWYLGIG--- ---------- ---------- ----DGGNIV RMHS------ 730
AV118_Gp37 LKTIKMICPV TINERNAA-- ---LTLD--- --SPSSSSAN YLQGSKAGTR SWYVGLG--- ---------- ---------- ----GSGNDL SLYS------ 737
AV112_Gp37 LKTIKMICPV TINERNAA-- ---LTLD--- --SPSSSSAN YLQGSKAGTR SWYVGLG--- ---------- ---------- ----GSGNDL SLYS------ 740
AV113_Gp37 LKTIKMICPV TINERNAA-- ---LTLD--- --SPSSSSAN YLQGSKAGTR SWYVGLG--- ---------- ---------- ----GSGNDL SLYS------ 740
AV117_Gp37 ----TSGQRI TVNTASDA-- ---IRIN--- --APTQESSN FIQGRKADVT KWYLGIG--- ---------- ---------- ----DGGNIV RMHS------ 727
   T4_Gp37 LTTINSNSRI NANFRMQLGQ SAYIDAECTD AVRPAGAGSF ASQNNEDVRA PFYMNIDRTD ASAYVPILKQ RYVQGNGCYS LGTLINNGNF RVHYHGGGDN 749
AV116_Gp37 LVVVTSNSRI SPNFRMQIGR SAYIDAECTD RVRPAGAGSF ASQNNENVRA PFYMNIDRTD ISTYVPIFKQ RYVQGNSCYS LGTLINGGNF RVHYHEGGDN 749
AV122_Gp37 LTTINSNSRI NANFRMQLGQ NAYIDIEASD GVRPTGCGSF ASQNDVNVRA PIYMNIERTA TSEYIPIVKQ RYVKGDSTYS IGTLISQGDF RVHYHQGTST 765
Conservation

AV110_Gp37 -------YTY SHGIALNSDT VDIAKPLKVG NAQFGTDGNI TGGTGNFANL NTTINNL--- ---------- ---------- ---------- ---------- 780
AV118_Gp37 -------QSY GHGLVISDNF VTISKPLKVG AAQLGTDGNI TGGSGNFANL NTTINNL--- ---------- ---------- ---------- ---------- 787
AV112_Gp37 -------QSY GHGLVISDNF VSITKPFKVG NAQLGTDGNI TGGSGNFANL NTTINNL--- ---------- ---------- ---------- ---------- 790
AV113_Gp37 -------QSY GHGLVISDNF VSITKPFKVG NAQLGTDGNI TGGSGNFANL NTTINNL--- ---------- ---------- ---------- ---------- 790
AV117_Gp37 -------YTY SHGIALNSDT VDIAKPLRIG STQIGTDGNI TGGSSNFANL NTTLERRVKV GTWSGSSTAG WYKFATVTMP QSVSTVQFKI IGGNGYNANS 820
   T4_Gp37 GSTGPQTADF GWEFIKNGDF IS-PRDLIAG KVRFDRTGNI TGGSGNFANL NSTIESL--- ---------- ---------- ---------- ---------- 805
AV116_Gp37 DSTGPLIKNF AWEFNRTGDF IS-PGKLGAG TVRIGTDGNI TGGSGNFANL NTTINNL--- ---------- ---------- ---------- ---------- 805
AV122_Gp37 N-----GVDL SWEFRRTGDF RA-PGKIIAG AATLGTDGNI TGGSGNFANL NTTIESL--- ---------- ---------- ---------- ---------- 816
Conservation

AV110_Gp37 ---------- ---------- ---------- ---------- ---------- ---------- ---------- ---------- ---------- ---------- 780
AV118_Gp37 ---------- ---------- ---------- ---------- ---------- ---------- ---------- ---------- ---------- ---------- 787
AV112_Gp37 ---------- ---------- ---------- ---------- ---------- ---------- ---------- ---------- ---------- ---------- 790
AV113_Gp37 ---------- ---------- ---------- ---------- ---------- ---------- ---------- ---------- ---------- ---------- 790
AV117_Gp37 PEQATITEVV LRTSNNNPKG INATLWGRTG NGFTNIATIN TSGDDYDVYL YAGTYADKLI VEYDATENTR YISVIGLNGG YQTPVTELPA GHLVGQVYMM 920
   T4_Gp37 ---------- ---------- ---------- ---------- ---------- ---------- ---------- ---------- ---------- ---------- 805
AV116_Gp37 ---------- ---------- ---------- ---------- ---------- ---------- ---------- ---------- ---------- ---------- 805
AV122_Gp37 ---------- ---------- ---------- ---------- ---------- ---------- ---------- ---------- ---------- ---------- 816
Conservation

AV110_Gp37 KTDIVSSYPI GAPIPWPTDT PPEGYAIMEG QTFDAGLYPK LAAVYPSGTL PDMRGQTIKG KPSGRAVLST EADGVKSHSH SASASSTDLG TKTTSSFDYG 880
AV118_Gp37 KTDIVSSYPI GAPIPWPTDT PPEGYAIMEG QTFDKSAYPK LAMAYPSGVI PDMRGQTIKG KPSGRAVLST EADGVKSHSH SASASSTDLG TKNTSSFDYG 887
AV112_Gp37 KTDIVSSYPI GAPIPWPTDT PPNGYAIMEG QTFDKSAYPK LAVAYPSGVI PDMRGQTIKG KPSGRAVLSA EADGVKSHSH SASASSTDLG TKTTSSFDYG 890
AV113_Gp37 KTDIVSSYPI GAPIPWPTDT PPNGYAIMEG QTFDKSAYPK LAVAYPSGVI PDMRGQTIKG KPSGRAVLSA EADGVKSHSH SASASSTDLG TKTTSSFDYG 890
AV117_Gp37 YSDAQIPYPV GAPIPWPSDS VPDGFALMEG QTFDKSAYPK LAIAYPTGTI PDMRGQTIKG KPSGRAVLSA EADGVKAHSH SASASSTDLG TKTTSSFDYG 1020
   T4_Gp37 KTDIMSSYPI GAPIPWPSDS VPAGFALMEG QTFDKSAYPK LAVAYPSGVI PDMRGQTIKG KPSGRAVLSA EADGVKAHSH SASASSTDLG TKTTSSFD-- 903
AV116_Gp37 KTDIVSSYPI GAPIPWPTDT PPEGYAIMEG QTFDKSAYPK LAMAYPSGVI PDMRGQTIKG KPSGRAVLST EADGVKSHSH SASASSTDLG TKNTSSFDYG 905
AV122_Gp37 KTDIVSSYPI GAPIPWPTDT PPDGYAIMEG QTFDKSAYPK LAMAYPSGVI PDMRGQTIKG KPSGRDVLSA EADGIKAHSH SASASSTDLG TKTTSSFDYG 916
Conservation

AV110_Gp37 SKTTSSFDYG TKTSNTTGNH NHTVSGNTSS AGAHQHARSG PQI-QAGIPN NIFYDGYNSV GANANAKFTG TVNGATAGSN IAKTSSDGAH THSWSGTTSA 979
AV118_Gp37 TKTSSSFDYG TKTSNTTGAH THSLSGSTNT TGAHTHSMQF VSGGTSGTPG ---------S GS-------- ----PDYSKY NANTSSAGAH SHSISGTAAS 966
AV112_Gp37 SKTTSSFDYG TKTSSTTGNH NHTVSGNTSS AGAHQHARSG PQI-QGGVPA SIFYDGYNSA GPNGNAKITG SVSGATASSN MAKTSSDGAH THSWSGTTSA 989
AV113_Gp37 SKTTSSFDYG TKTSSTTGNH NHTVSGNTSS AGAHQHARSG PQI-QGGVPA SIFYDGYNSA GPNGNAKITG SVSGATASSN MAKTSSDGAH THSWSGTTSA 989
AV117_Gp37 TKTTSSFDYG TKTSNNTGAH THSLSGSTNT AGAHSHTSAL YAGD-RNSPG G--------- ---------S SARGSAGGGN TLGTSSSGAH SHSISGTAAS 1101
   T4_Gp37 --------YG TKGTNSTGGH THSGSGSTST NGEHSHYIEA WNGTGV---- ---------- --GGNKMS-- -SYAISYRAG GSNTNAAGNH SHTFSFGTSS 976
AV116_Gp37 TKTSSSFDYG TKTSNTTGAH THSLSGSTNT TGAHTHSMQF VSGGTS---- ---------- --GTPGSG-- ---SPDYSKY NANTSSAGAH SHSISGTAAS 984
AV122_Gp37 TKTSSSFDYG TKNTNSTGGH THSVSGTTSA AGEHAHHLGL LLVNGGDA-- ---LYGYTTV GNSLPRTL-- -DWLDRRDNK FPNTNTTGNH THTWSGTTSN 1008
Conservation

AV110_Gp37 TGNHAHTVGI GAHTHSVGIG AHSHTVAIGS HGHTITVNAT GNTENTVKNV AFNYIVRLA- 1038
AV118_Gp37 AGAHAHTVGI GAHTHTVGIG AHTHTVAIGS HGHTITVNNT GNTENTVKNI AFNYIVRLA- 1025
AV112_Gp37 TGNHAHTVGI GAHTHSVGIG AHTHTVAIGS HGHTITVNAT GNTENTVKNV AFNYIVRLA- 1048
AV113_Gp37 TGNHAHTVGI GAHTHSVGIG AHTHTVAIGS HGHTITVNAT GNTENTVKNV AFNYIVRLA- 1048
AV117_Gp37 AGAHAHTVGI GAHTHTVGIG AHTHSVAIGS HGHTITVNST GNTENTVKNI AFNYIVRLA- 1160
   T4_Gp37 ---------A GDHSHSVGIG AHTHTVAIGS HGHTITVNST GNTENTVKNI AFNYIVRLA- 1026
AV116_Gp37 AGAHAHTVGI GAHTHTVGIG AHTHTVAIGS HGHTITVNNT GNTENTVKNI AFNYIVRLA- 1043
AV122_Gp37 AGAHTHTVGI GAHTHTVGIG AHTHSVAIGS HGHTITVNST GNTENTVKNI AFNYIVRLA- 1067
Conservation
```
